# Transit-amplifying cells coordinate changes in intestinal epithelial cell-type composition

**DOI:** 10.1101/840371

**Authors:** Laura E. Sanman, Ina W. Chen, Jake M. Bieber, Veronica Steri, Byron Hann, Lani F. Wu, Steven J. Altschuler

## Abstract

Renewing tissues have the remarkable ability to continually produce both proliferative progenitor and specialized differentiated cell-types. How are complex milieus of microenvironmental signals interpreted to coordinate tissue cell-type composition? Here, we develop a high-throughput approach that combines organoid technology and quantitative imaging to address this question in the context of the intestinal epithelium. Using this approach, we comprehensively survey enteroid responses to individual and paired perturbations to eight epithelial signaling pathways. We uncover culture conditions that enrich for specific cell-types, including Lgr5^+^ stem and enteroendocrine cells. We analyze interactions between perturbations and dissect mechanisms underlying an unexpected mutual antagonism between EGFR and IL-4 signals. Finally, we show that, across diverse perturbations, modulating proliferation of transit-amplifying cells also consistently changes the composition of differentiated secretory and absorptive cell-types. This property is conserved *in vivo* and can arise from differential amplification of secretory and absorptive progenitor cells. Taken together, the observations highlight an underappreciated role for transit-amplifying cells in which proliferation of these short-lived progenitors provides a lineage-based mechanism for tuning differentiated cell-type composition.

## Introduction

A central question in the study of complex tissues is how diverse signals are integrated to regulate cell-type composition. Dissection of mechanisms underlying the mapping from signals to tissue composition is complicated by the heterogeneous makeup of interconnected cell-types, which exert influences upon one another through the lineage structure and cell-cell interactions. Further, most studies focus on the effects of individual signals on individual cell-types, due to a lack of methods that enable systematic identification of signal integration mechanisms. What are the tissue-wide effects of common microenvironmental signals on cell-type composition? How do multiple signals modify each other’s effects? Finally, are there intrinsic tissue properties that shape response to diverse signals?

Here, we address these questions in the context of the intestinal epithelium, a model renewing tissue (Beumer & Clevers, 2016; Cheng & Leblond, 1974b; A. Tian et al., 2016). The intestinal epithelium is particularly remarkable in that it maintains a stereotypic tissue composition despite a rapid 3- to 5-day turnover. During renewal of the intestinal epithelium, Lgr5^+^ crypt-base stem cells differentiate into proliferating transit-amplifying (TA) progenitors, which in turn adopt absorptive (enterocyte) or secretory (Paneth, goblet, enteroendocrine (EE)) cell fates (Cheng & Leblond, 1974b) (Figure 1A). The confluence of proliferating and differentiation decision processes establishes tissue composition, which guides overall tissue function. Though much progress has been made in defining the ‘parts list’ of factors that guide intestinal epithelial renewal (Clevers, 2013; Flier et al., 2009; Yin et al., 2014a), it is unclear how these factors are integrated by the tissue during maintenance and in response to perturbations.

**Figure 1.**
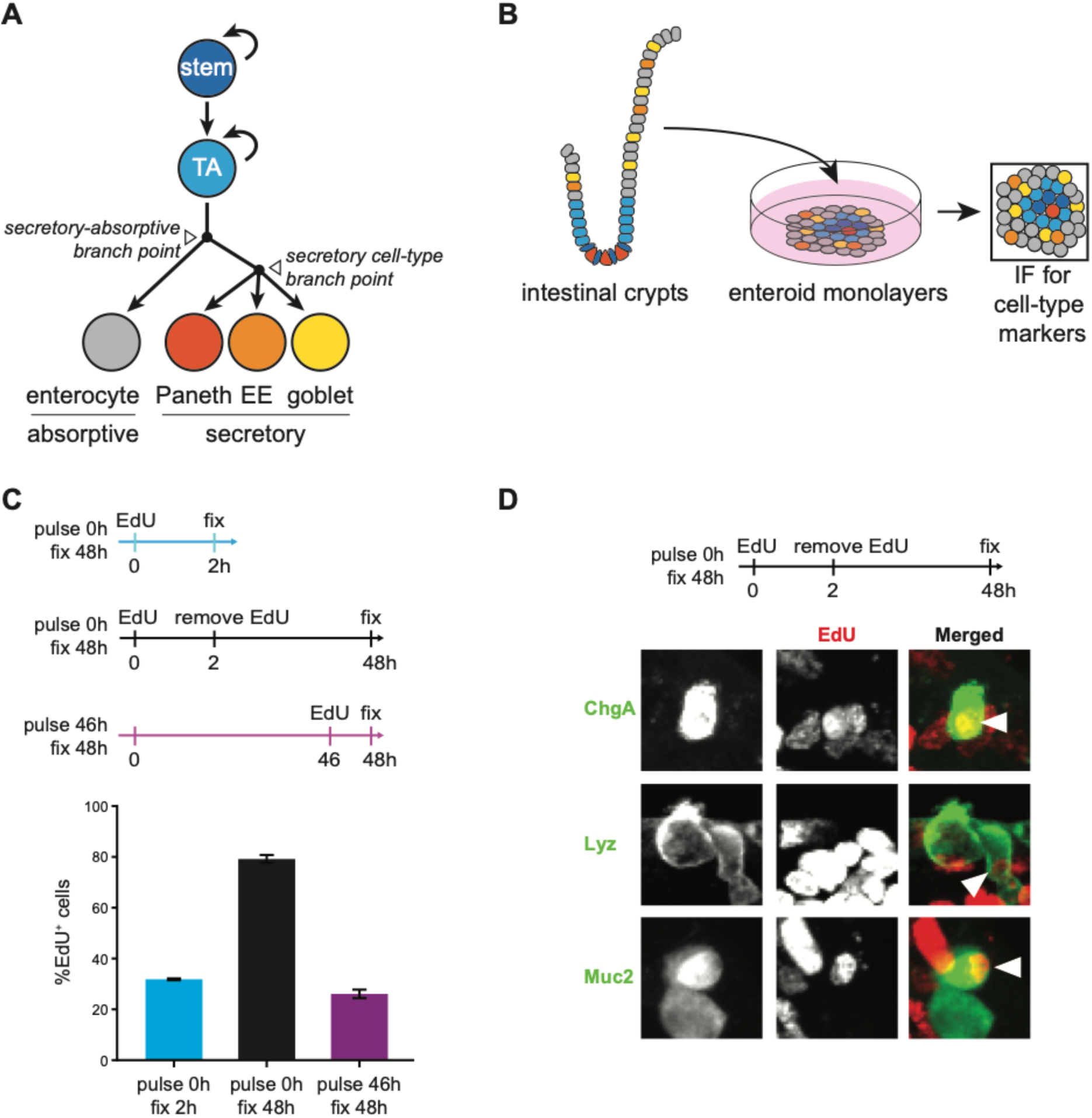
A quantitative microscopy platform to study regulation of intestinal cell-type composition. (**A**) Intestinal epithelial cell lineage diagram. (**B**) Workflow for characterization of perturbation effects on cell-type composition. Intestinal crypts are cultured as enteroid monolayers. Cell-type numbers are measured from immunofluorescence (IF) images of enteroid monolayers. (**C**) ∼80% of cells in 48 hours of enteroid culture are produced by initial progenitors. Enteroid monolayers were seeded and either EdU was added for 2 hours and enteroids were then fixed (cyan), EdU was pulsed for 2 hours and removed for 46 hours before fixation (black), or enteroids were cultured for 46 hours and EdU added for the 2 hours prior to fixation (magenta). ∼20-30% of cells are proliferating (EdU^+^) upon initial seeding (cyan) and after 48 hours of culture (magenta), and ∼80% of cells are produced by initially proliferating progenitors (EdU^+^) in the 48-hour culture period (black). n=3 wells. (**D**) Initial progenitors produce secretory cell-types during 48 hours of culture. Enteroids were pulsed with EdU for 2 hours upon initial seeding. EdU was removed and cells were fixed after 46 hours of culture. EdU and cell-type marker (Lyz; Paneth, Muc2; goblet, and ChgA; EE) staining patterns were then visualized. Arrowheads indicate cells that co-stain for EdU and the indicated cell-type markers. Error bars mean +/− sem.

To quantitatively measure intestinal epithelial cell-type composition and study its changes in response to microenvironmental signals, we utilized a recently established enteroid monolayer culture system that recapitulates key features of the intestinal epithelium (Thorne et al., 2018). Enteroid monolayer cultures maintain characteristics of intestinal epithelial architecture, including organization into crypt- and villus-like compartments and apical-basolateral polarization. Importantly, these cultures also preserve core tissue processes *ex vivo*, generating all major intestinal epithelial cell-types (Lgr5^+^ stem, TA, and differentiated secretory and absorptive cells) with a turn-over rate similar to the *in vivo* renewal rate. Crucially, due to their 2D nature, enteroid monolayer cultures are amenable to high-throughput screening of perturbations in microwell format. In this work, enteroid monolayers are used as a primary platform for hypothesis generation with key observations further evaluated in 3D organoids and *in vivo*.

Here, we present a systems approach for investigating signal integration and lineage processes in the intestinal epithelium. We first expanded the capabilities of the enteroid monolayer platform to monitor and quantify major intestinal epithelial cell-types, such as stem, TA, and differentiated cell-types, in a robust and automated manner. This enabled confirmation of relevant stem and TA functions, such as proliferation and production of differentiated cell-types, in enteroid monolayer cultures. We then combined enteroid culture with high-throughput combinatorial perturbations to profile responses to a diverse set of treatment conditions. Analysis of tissue response profiles identified conditions that enrich for specific intestinal epithelial cell-types and revealed interactions in the integration of perturbations. Finally, studies using enteroid monolayers, 3D organoids, and mice uncovered a global trend in how intestinal epithelial cell-type composition changes in response to perturbations; specifically, a link between TA proliferation and secretory cell composition. Based on experimental observations and mathematical modeling, we propose a model of intestinal epithelial lineage control in which differential amplification of TA progenitors allows the balance of secretory to absorptive cell lineages to be regulated through TA proliferation.

## Results

### A quantitative microscopy platform to study regulation of intestinal cell-type composition

As an experimental model for the study of intestinal epithelial cell-type composition in high-throughput, enteroid monolayers should be quantifiable, reproducible, and faithfully recapitulate relevant *in vivo* tissue properties. Here, we develop and test these features of the enteroid monolayer platform.

To measure intestinal epithelial composition in high-throughput, we established a computational pipeline to automatically quantify major intestinal cell-types from images of enteroid monolayers (Figure 1B). Specifically, we developed algorithms to detect cells expressing markers for stem (Lgr5^+^), proliferating (EdU^+^), Paneth (Lyz^+^), goblet (Muc2^+^), and enteroendocrine (EE; ChgA^+^) cells as well as identifying cell nuclei (Hoechst) (Figure 1-Supplement 1, Materials and Methods). Enterocytes were not directly quantified because we were not able to identify a reliable antibody to specifically label this population in immunofluorescence assays. Thus, our studies of differentiated cell-types were limited to the secretory lineage. When evaluated against expert manual counting, the algorithms exhibited high quantification accuracy across the cell-types measured (Table 1).

**Table 1.** Evaluation of the performance of cell-type identification algorithms.

| Cell-type | Marker | Precision | Recall | F1 Score |
| --- | --- | --- | --- | --- |
| nuclei | Hoechst | 0.9965 | 0.9936 | 0.9951 |
| goblet cells | Muc2 | 1 | 0.8998 | 0.9473 |
| EE cells | ChgA | 0.8551 | 0.9248 | 0.8886 |
| Paneth cells | Lyz | 0.9018 | 0.9642 | 0.9319 |
| EdU+ cells | EdU | 1 | 0.9557 | 0.9774 |
| crypt regions | Lgr5-GFP | 0.9816 | 0.9877 | 0.9846 |
| stem cells | Lgr5-GFP | 0.9724 | 0.9756 | 0.9740 |
| Lgr5+/Dclk1+ cells | Lgr5-GFP | 0.8208 | 0.9087 | 0.8625 |

To evaluate reproducibility, cell-type measurements were quantified across replicate wells. A crypt seeding density (10-20% initial confluency) was identified where there was no relationship between initial seeding density and cell-type composition after 48 hours of culture. In this regime, there was also relatively low inter-replicate variability in cell-type numbers. (Figure 1-Supplement 2).

To assess recapitulation of relevant *in vivo* intestinal epithelial properties, we examined cell-type composition and progenitor cell functions including proliferation and differentiation. We first observed that jejunal enteroid monolayers exhibited cell-type composition comparable to the composition of *in vivo* jejunal epithelium (Table 2). We then confirmed that enteroid monolayers recapitulate progenitor proliferation and production of differentiated cell-types. By labeling cycling cells with EdU upon initial plating, we observed that the initial population of ∼25% proliferative (EdU^+^) cells produce the vast majority (∼80%) of cells in enteroid monolayers after 48 hours of culture (Figure 1C). This indicated that the majority of quantified cells after 48 hours of culture were produced by initial progenitors. Further, we observed EdU^+^ cells that colocalized with markers of Paneth (Lyz), goblet (Muc2), and EE (ChgA) cells, indicating that progenitor cells produce major differentiated cell-types over the 48-hour culture period (Figure 1D).

**Table 2.**
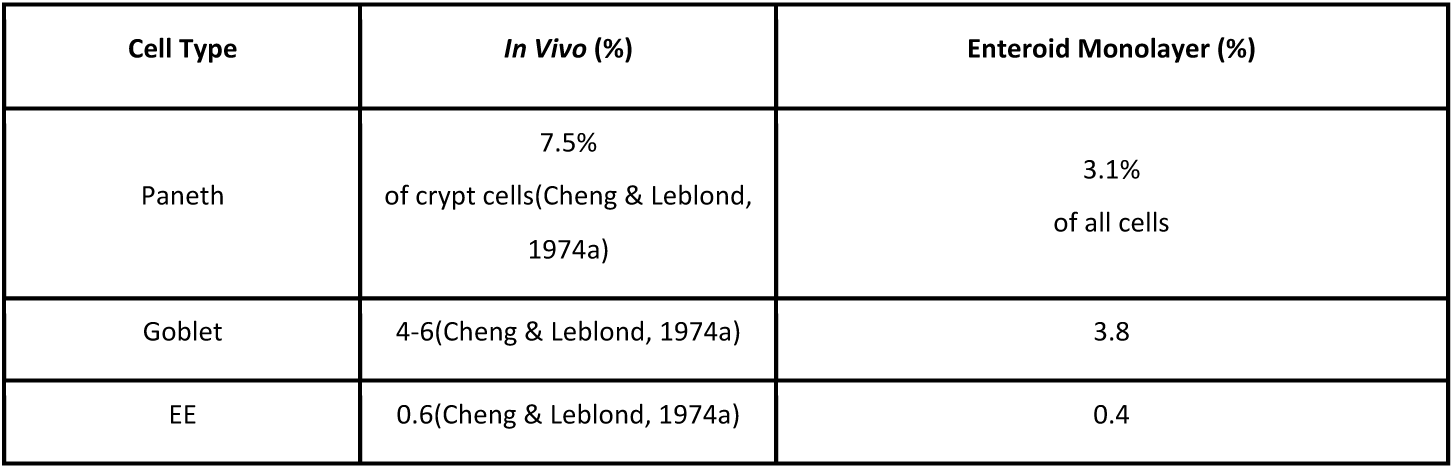
Comparison of enteroid monolayer cell-type composition with literature reports of small intestine cell-type composition.

| Cell Type | *In Vivo* (%) | Enteroid Monolayer (%) |
| --- | --- | --- |
| Paneth | 7.5%<br>of crypt cells(Cheng & Leblond,<br>1974a) | 3.1%<br>of all cells |
| Goblet | 4-6(Cheng & Leblond, 1974a) | 3.8 |
| EE | 0.6(Cheng & Leblond, 1974a) | 0.4 |

Taken together, enteroid monolayers preserve important characteristics of the intestinal epithelium *ex vivo*, such as cell-type composition and production of differentiated cell-types. Enteroid monolayers also demonstrate reproducible outgrowth and low replicate variance. Thus, enteroid monolayers constitute a robust platform for capturing tissue-wide responses to perturbations and for generating hypotheses on control of intestinal epithelial cell-type composition.

### Systematic survey of cell-type composition changes in response to single and pairwise signaling modulators

To map a wide range of tissue composition phenotypes, 13 epithelial-intrinsic and microenvironmental modulators that target eight core intestinal epithelial signaling pathways (Wnt, BMP, Notch, HDAC, JAK, p38MAPK, TGF-beta, EGFR (Basak et al., 2017a; Batlle et al., 2002; Mathieu Houde et al., 2001; Richmond et al., 2018a; Rodríguez-Colman et al., 2017; van der Flier & Clevers, 2009; Yin et al., 2014b)) and are known to have diverse effects on tissue cell-type composition were selected (Table 3). In previous reports, combinations of perturbations have been shown to be more effective than single perturbations in enriching particular cell-types (Basak et al., 2017a; Yin et al., 2014b). Thus, to survey a broad range of tissue states, modulators were applied to enteroid monolayers individually (13 conditions) and in all possible pairwise combinations (78 conditions) (workflow: Figure 2-Supplement 1A). Perturbation concentrations were guided by literature reports and confirmed with dose-response experiments (Table 3, Figure 2-Supplement 1B) (Basak et al., 2017a; von Moltke et al., 2015; Yin et al., 2014b).

**Table 3.**
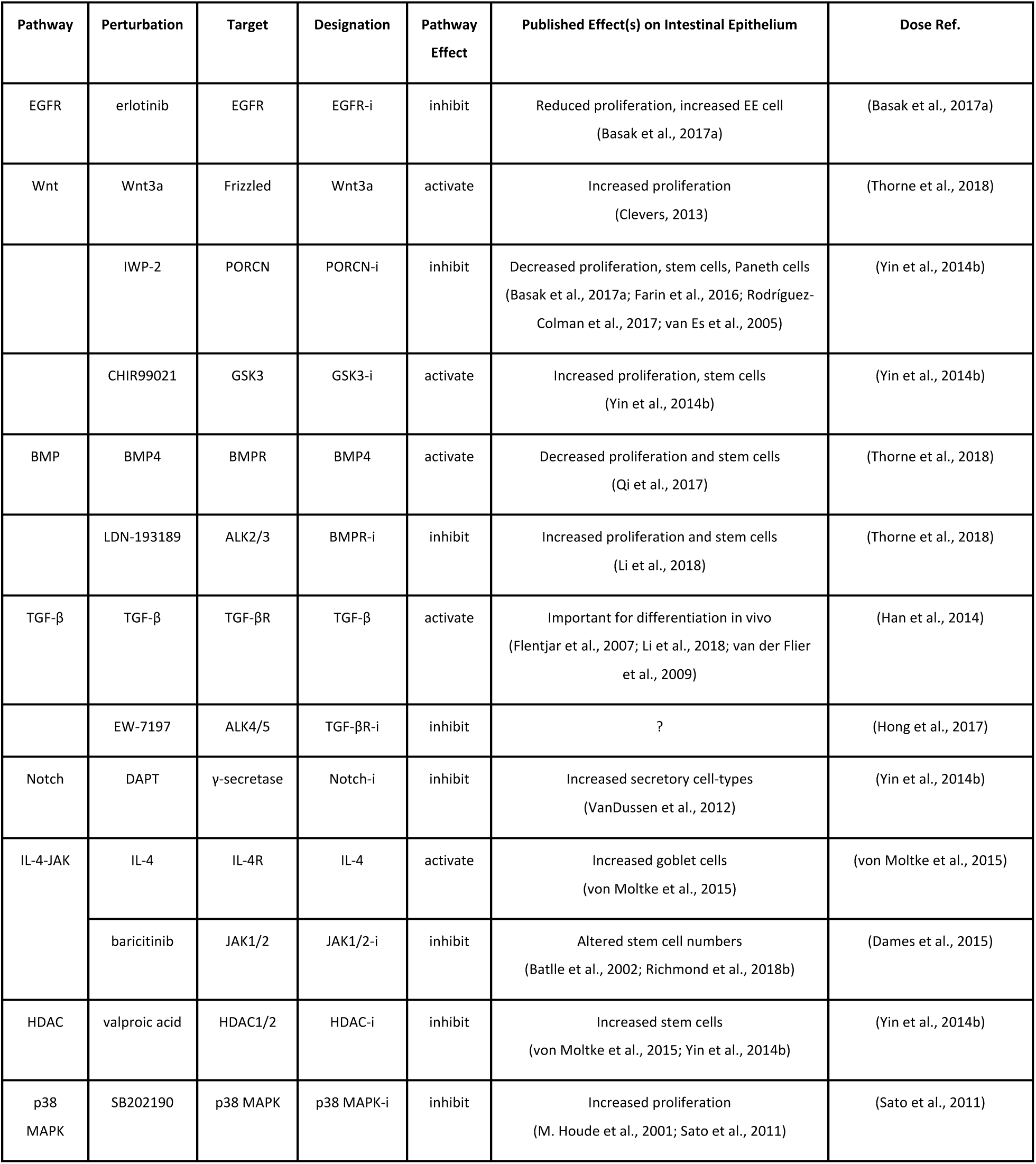
Selection of cell-type perturbations for study. For each perturbation, the specific pathway and target are indicated, as is the expected effect on the pathway and tissue. Designation is the code with which the perturbation is referred to in figures and text.

| Pathway | Perturbation | Target | Designation | Pathway Effect | Published Effect(s) on Intestinal Epithelium | Dose Ref. |
| --- | --- | --- | --- | --- | --- | --- |
| EGFR | erlotinib | EGFR | EGFR-i | inhibit | Reduced proliferation, increased EE cell<br>(Basak et al., 2017a) | (Basak et al., 2017a) |
| Wnt | Wnt3a | Frizzled | Wnt3a | activate | Increased proliferation<br>(Clevers, 2013) | (Thorne et al., 2018) |
|  | IWP-2 | PORCN | PORCN-i | inhibit | Decreased proliferation, stem cells, Paneth cells<br>(Basak et al., 2017a; Farin et al., 2016; Rodríguez-Colman et al., 2017; van Es et al., 2005) | (Yin et al., 2014b) |
|  | CHIR99021 | GSK3 | GSK3-i | activate | Increased proliferation, stem cells<br>(Yin et al., 2014b) | (Yin et al., 2014b) |
| BMP | BMP4 | BMPR | BMP4 | activate | Decreased proliferation and stem cells<br>(Qi et al., 2017) | (Thorne et al., 2018) |
|  | LDN-193189 | ALK2/3 | BMPR-i | inhibit | Increased proliferation and stem cells<br>(Li et al., 2018) | (Thorne et al., 2018) |
| TGF-β | TGF-β | TGF-βR | TGF-β | activate | Important for differentiation in vivo<br>(Flentjar et al., 2007; Li et al., 2018; van der Flier et al., 2009) | (Han et al., 2014) |
| | EW-7197 | ALK4/5 | TGF-βR-i | inhibit | ? | (Hong et al., 2017) |
| Notch | DAPT | γ-secretase | Notch-i | inhibit | Increased secretory cell-types<br>(VanDussen et al., 2012) | (Yin et al., 2014b) |
| IL-4-JAK | IL-4 | IL-4R | IL-4 | activate | Increased goblet cells<br>(von Moltke et al., 2015) | (von Moltke et al., 2015) |
|  | baricitinib | JAK1/2 | JAK1/2-i | inhibit | Altered stem cell numbers<br>(Battle et al., 2002; Richmond et al., 2018b) | (Dames et al., 2015) |
| HDAC | valproic acid | HDAC1/2 | HDAC-i | inhibit | Increased stem cells<br>(von Moltke et al., 2015; Yin et al., 2014b) | (Yin et al., 2014b) |
| p38 MAPK | SB202190 | p38 MAPK | p38 MAPK-i | inhibit | Increased proliferation<br>(M. Houde et al., 2001; Sato et al., 2011) | (Sato et al., 2011) |

We then sought to combine cell-type number measurements into readouts that reflect changes in important aspects of intestinal epithelial cell-type composition. Changes in overall tissue composition can be attributed to a number of changes in underlying cell-type populations. For instance, an overall decrease in the fraction of a specific cell-type, such as goblet cells, may be caused by an increase in the number of progenitors (or other cell-types), a decrease in the secretory fraction in the differentiated cell population, or a decrease in goblet cells within the secretory cell lineage. To distinguish between these different possibilities, we designed features that measured the size of the progenitor population as well as the relative composition of cells in the differentiated population, as follows.

To quantify changes in the progenitor cell populations, we measured the absolute numbers of proliferating stem (EdU^+^ Lgr5^+^) and TA (EdU^+^ Lgr5^-^) cells (Figure 2A, Figure 2-Supplement 2A). Since specific TA markers are lacking, TA cells were defined as non-stem proliferating cells. To quantify changes in the relative makeup of the differentiated population, we designed fractional readouts that reflect changes in two differentiation branching points: the secretory-absorptive branch point, and the secretory cell-type branch point. To capture secretory vs. absorptive lineage biases, we computed the fraction of differentiated (Lgr5^-^ EdU^-^) cells that express secretory (Paneth+goblet+EE) markers (#secretory/#diff) (Figure 2A, Figure 2-Supplement 2B). The sum of Paneth, goblet, and EE cells is a reasonable approximation for the number of secretory cells as these cell-types make up the majority of secretory cells (Cheng & Leblond, 1974b; Haber et al., 2017). Finally, to capture biases towards different cell-types within the secretory lineages, we computed the specific fractions of all secretory cells positive for Paneth (Lyz), goblet (Muc2), and EE (ChgA) cell markers (#Paneth/#secretory, #goblet/#secretory, #EE/#secretory) (Figure 2A, Figure 2-Supplement 2C).

**Figure 2.**
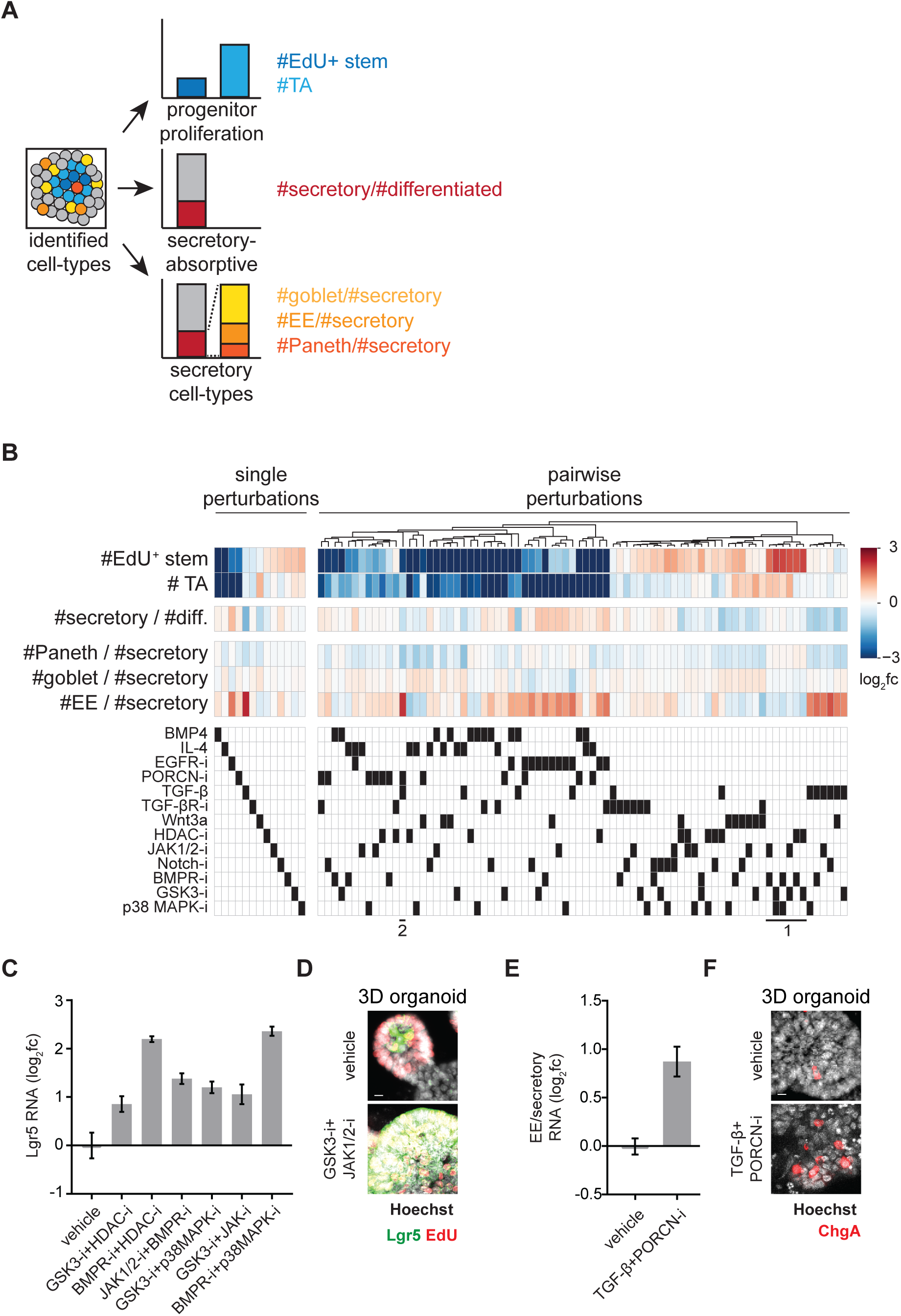
Systematic characterization of perturbation effects on intestinal epithelial cell-type composition reveals cell-type-specific regulators. (**A**), Cell-type counts are combined to quantify progenitor proliferation (top), overall secretory cell prevalence (middle), and representation of each secretory cell-type within that lineage (bottom). (**B**) Combinatorial perturbations induce diverse phenotypes. Single (left) and pairwise (right) perturbation effects are shown. Top colored heatmaps: Perturbation effects are represented as log_2_ fold-change (fc) relative to vehicle-treated wells. Bottom black-and-white heatmap: Black boxes indicate perturbations in each column. Single perturbations are sorted by #EdU^+^ stem cells; combinatorial perturbations are clustered based on similarity of tissue-wide effects. Callouts (1) and (2) referred to in text. n=2-28 wells. (**C-D**) Recapitulation of Lgr5^+^ stem cell-inducing conditions by qRT-PCR in enteroid monolayers and IF in 3D organoids. (**C**) Enteroid monolayers were treated as indicated for 48 hours. RNA levels were measured by qRT-PCR. n=3 wells. (**D**) Lgr5-GFP-DTR 3D organoids were treated as indicated for 48 hours. EdU was added for 2 hours prior to fixation. (**E-F**) Recapitulation of EE cell inducing condition by qRT-PCR in enteroid monolayers and IF in 3D organoids. (**E**) Enteroid monolayers were treated as indicated for 48 hours. Changes in EE cell RNA (ChgA) relative to average of secretory cell RNA (ChgA+Lyz+Muc2) was measured by qRT-PCR. n=3 wells. (**F**) 3D organoids were treated as indicated for 24 hours and increased EE (ChgA^+^) cell numbers were observed. Error bars mean +/− sem.

After treatment with established modulators of cell-type composition (GSK3-i+HDAC-i, Notch-i, IL-4, EGFR-i, PORCN-i), changes in enteroid monolayer cell-type readouts were in agreement with previous 3D organoid and *in vivo* studies (Basak et al., 2017a; Beumer & Clevers, 2016; Qi et al., 2017; van Es et al., 2005; von Moltke et al., 2015; Yin et al., 2014b) (Figure 2-Supplement 3A, Table 3). Changes in the prevalence of each cell-type were also confirmed at the RNA level using quantitative reverse transcription PCR (qRT-PCR; Figure 2-Supplement 3B).

### Identification of conditions that enrich for Lgr5**^+^** stem and EE cells

Single and pairwise perturbation effects on intestinal epithelial cell-type composition (546 measurements = 91 conditions x 6 cell-type readouts) were measured (Figure 2B). The vast majority of measurements were previously uncharacterized, with a number of conditions strongly modulating different aspects of tissue composition (Table 4). Below, we highlight conditions that were previously unknown to enrich for stem and EE cells.

**Table 4.** Top perturbations in modulating cell-type composition.

|  | Downregulating |  | Upregulating |  |
| --- | --- | --- | --- | --- |
|  | Perturbation | Log<sub>2</sub>fc | Perturbation | Log<sub>2</sub>fc |
| EdU+ stem number | BMP4 + JAK1/2-i | -11.380 | GSK3-i + p38 MAPK-i | 2.602 |
|  | BMP4 | -11.080 | GSK3-i + JAK1/2-i | 2.405 |
|  | BMP4 + Wnt3a | -10.380 | BMPR-i + p38 MAPK-i | 2.363 |
| TA number | EGFR-i + HDAC-i | -9.476 | GSK3-i + Wnt3a | 1.541 |
|  | IL-4 + PORCN-i | -9.437 | GSK3-i + JAK1/2-i | 1.415 |
| | EGFR-i + TGF-β-i | -8.512 | BMPR-i + Wnt3a | 1.240 |
| Secretory fraction<br>(of differentiated cells) | TGFβ + Wnt3a | -1.544 | GSK3-i + EGFR-i | 1.109 |
| | BMP4 + TGFβ | -1.509 | EGFR-i + BMPR-i | 1.096 |
| | TGFβ + HDAC-i | -1.452 | EGFR-i + JAK1/2-i | 1.062 |
| Paneth fraction<br>(of secretory cells) | IL-4 + TGF-β | -1.651 | BMPR-i + HDAC-i | 0.307 |
| | PORCN-i + TGF-β | -1.466 | GSK3-i + HDAC-i | 0.305 |
| | IL-4 + p38 MAPK-i | -1.417 | EGFR-i + TGF-β | 0.270 |
| Goblet fraction<br>(of secretory cells) | EGFR-i + TGF-β | -1.209 | IL-4 + p38 MAPK-i | 0.724 |
| | Notch-i + EGFR-i | -0.880 | TGF-β-i + Wnt3a | 0.611 |
| | EGFR-i + PORCN-i | -0.558 | IL-4 + TGF-β | 0.604 |
| EE fraction<br>(of secretory cells) | TGF-β-i + Wnt3a | -1.525 | PORCN-i + TGF-β | 2.885 |
| | IL-4 + HDAC-i | -1.419 | Notch-i + TGF-β | 2.542 |
| | GSK3-i + HDAC-i | -1.098 | TGF-β | 2.282 |

Pairwise combinations of inhibitors of GSK3, p38MAPK, BMPR, HDAC, and JAK1/2 increased the number of Lgr5^+^ stem cells (Figure 2B, bottom callout 1). Notably, these conditions caused similar, if not increased, enrichment for Lgr5^+^ stem cells compared to the current benchmark condition (GSK3 inhibitor + HDAC inhibitor (Yin et al., 2014a)). Lgr5^+^ stem cell enrichment was also observed at the RNA level, as indicated by qRT-PCR analysis of Lgr5 RNA (Figure 2C). Since JAK1/2 had not been connected with stemness in the mammalian intestinal epithelium in the absence of inflammation (Richmond et al., 2018b), we repeated the GSK3-i and JAK1/2-i treatment in 3D organoids and again observed expansion of the Lgr5^+^ stem cell population (Figure 2D).

Conditions were also identified that selectively enrich for EE cells relative to other secretory cell-types (Table 4). The condition that most strongly enriched for EE cells was the combination of TGF-beta and PORCN inhibitor (Figure 2B, bottom callout 2). TGF-beta and PORCN inhibitor treatment of enteroid monolayers also caused an increase in RNA levels of the EE marker ChgA relative to RNA levels of markers of other secretory cell-types (Figure 2E). Since a connection between TGF-beta signaling and EE cell regulation had not been previously established, we applied the treatment to 3D organoids and again observed EE enrichment (Figure 2F).

### Interaction mapping reveals mutual antagonism between IL-4 and EGFR-i

After analyzing perturbations effects individually, we next examined how pairs of perturbations interact to modulate cell-type composition. A commonly used framework for identifying perturbation interactions is a multiplicative model, where the combined effect of two non-interacting perturbations is expected to be the product of the effects of individual perturbations (Dixon et al., 2009). For each cell-type composition readout, we analytically defined an interaction between any pair of perturbations as a combinatorial perturbation effect that deviated strongly (effect size>5) from the expectation of the multiplicative model. (The multiplicative model was chosen due to its interpretability and fit to the data; Figure 3-Supplement 1, Materials and Methods). Biologically, interactions were interpreted as direct or indirect connections between the perturbed signaling pathways (Coster et al., 2017; Papin et al., 2005). The majority of perturbation pairs showed little deviation from the multiplicative model expectation (Figure 3-Supplement 1A). Interestingly, perturbation interactions were predominantly observed in the regulation of EdU^+^ stem and TA cell numbers, as compared to the regulation of secretory, goblet, Paneth, or EE cell prevalence (Figure 3A).

**Figure 3.**
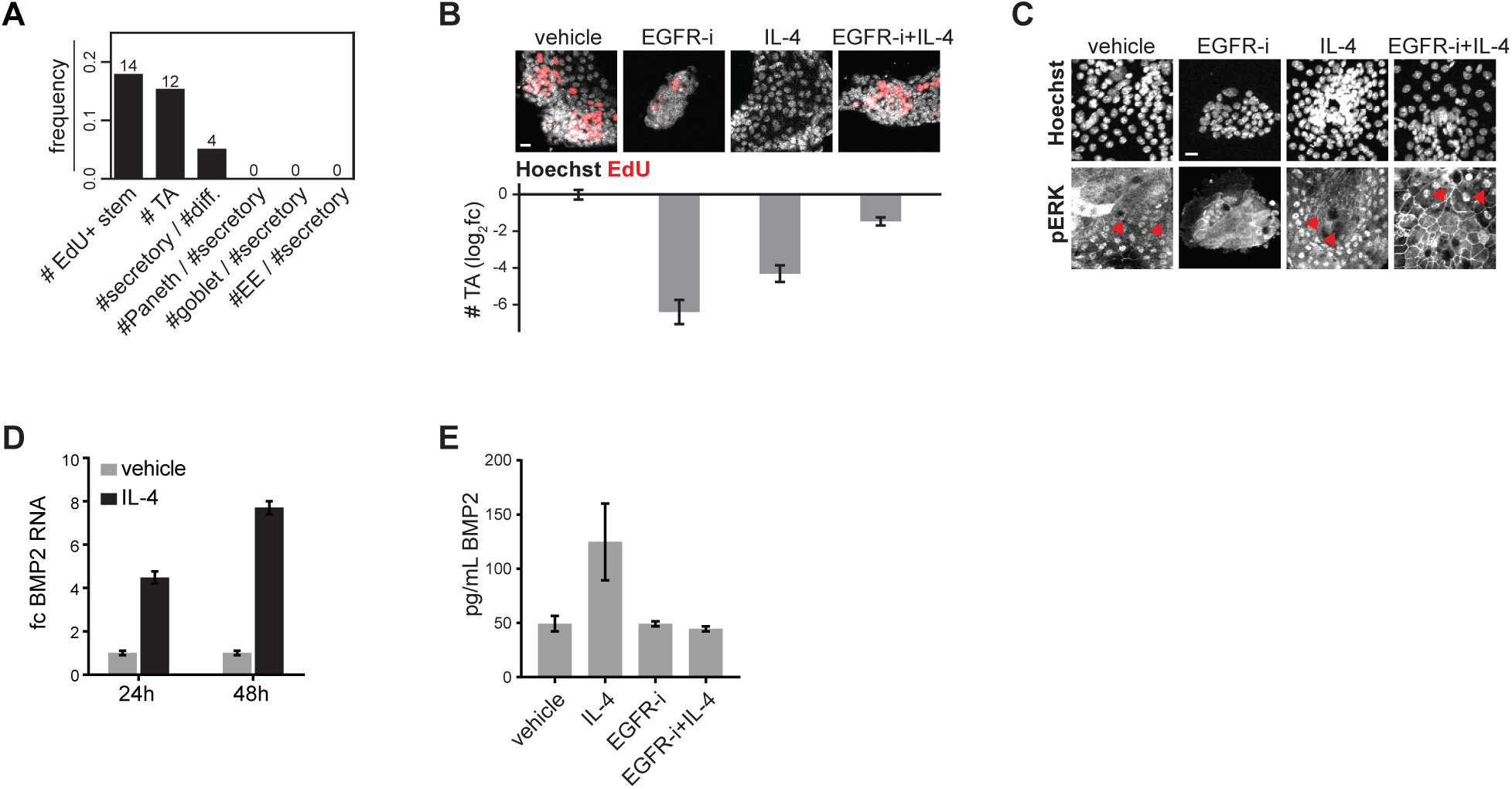
Interaction mapping reveals mutual antagonism between EGFR-i and IL-4. (**A**) Perturbation interactions are predominantly observed in the regulation of EdU^+^ stem and TA cell numbers. The frequency of perturbation pairs that exhibit significant deviation from the multiplicative expectation was quantified for each readout of cell-type composition. (**B**) EGFR-i and IL-4 antagonize each other’s effects on TA numbers. Enteroid monolayers were treated as indicated for 48 hours. Images (top) and quantification of TA cell numbers (bottom) are shown. n=2-6 wells. (**C**) IL-4 induces phospho-Erk nuclear translocation in the context of EGFR inhibition. Enteroid monolayers were treated as indicated for 48 hours and then stained for phospho-Erk. Nuclear phospho-Erk is observed in all conditions except EGFR-i alone. Red arrowheads indicate example cells with nuclear phospho-Erk. (**D**) IL-4 induces BMP2 production. Enteroid monolayers were treated as indicated and BMP2 RNA levels measured by qRT-PCR. n=3 wells. (**E**) EGFR-i abrogates the IL-4-induced increase in BMP2 production. Enteroid monolayers were treated as indicated for 48 hours and levels of BMP2 in the media measured by ELISA. n=3 wells. Error bars mean +/− sem. Fc=fold-change.

In some cases, identified interactions reflected known pathway biology, such as interactions between perturbations that modulate the same signaling pathway (e.g. the synergistic effect of Wnt3a and GSK3-i, Figure 3-Supplement 2). Other cases were less expected and suggest novel interactions in the regulation of specific cell-types. One particularly unexpected example is the interaction between IL-4 and EGFR-i in regulating TA cells. IL-4 and EGFR-i are not known to interact or utilize a shared signaling pathway and, interestingly, appeared mutually antagonistic in combination. When either IL-4 or EGFR-i was applied singly, the number of TA cells was reduced by ∼4 and ∼6 log-fold relative to control, respectively. However, when IL-4 and EGFR-i were applied in combination, the number of TA cells was reduced by only ∼1.5 log-fold relative to control (Figure 3B).

We investigated potential mechanisms for the observed mutual antagonism of IL-4 and EGFR-i. EGFR-i treatment decreases proliferation by reducing MEK-Erk activity (Basak et al., 2017a). However, IL-4 treatment resulted in Erk activation (as measured by phospho-Erk nuclear translocation) even in the context of EGFR inhibition (Figure 3C). This suggested that IL-4 antagonized the effects of EGFR-i by bypassing EGFR-i to activate Erk. When IL-4 and EGFR-i were applied in the context of MEK inhibition (which is upstream of Erk), IL-4 did not antagonize the effects of EGFR-i on TA cell numbers (Figure 3-Supplement 3A) or induce phospho-Erk nuclear translocation (Figure 3-Supplement 3B). This indicated that IL-4 activates MEK-Erk signaling upstream of MEK. In the other direction, IL-4 reduced proliferation in both enteroid monolayers and 3D organoids by increasing BMP2 (the epithelial paralog of mesenchymal BMP4 (Goldman et al., 2009)) production by the tissue (Figure 3D). The addition of a BMP receptor inhibitor, BMPR-i, rescued the IL-4-induced decrease in transit-amplifying cell numbers further confirming BMP secretion as the mechanism by which IL-4 dampens proliferation (Figure 3-Supplement 4A,B). EGFR-i reduces BMP2 production induced by IL-4 treatment, suggesting a mechanism by which EGFR-i antagonizes the effects of IL-4 (Figure 3E).

In summary, analysis of perturbation interactions, which occurred predominantly in progenitor cells, revealed a surprising mutual antagonism between IL-4 and EGFR signaling in the regulation of TA cells. Mechanistic investigation revealed that the mutual antagonism arises from IL-4 bypassing EGFR-i-induced blockade of MEK-Erk-driven cell cycle progression and EGFR-i blocking IL-4-induced production of antiproliferative BMP2.

### Modulating transit-amplifying cell proliferation alters secretory cell-type composition

In addition to gaining insights into the signaling regulation of tissue composition, we applied this dataset to survey global trends in how intestinal epithelial cell-type composition changes in response to perturbations. In particular, the observation of increased interaction in regulation of the progenitor populations led us to investigate how changes in progenitor cell numbers relate to changes in the composition of differentiated cell-types. Despite the diversity of perturbation mechanisms, we observed a surprising trend: the numbers of transit-amplifying cells negatively correlated with the fraction of differentiated cells that express secretory markers (Figure 4A) as well as with the fraction of secretory cells that express the EE marker (Figure 4-Supplement 1A). In contrast, little-to-no correlation was observed between the number of proliferating Lgr5^+^ stem cells and differentiated cell-type composition (Figure 4A, Figure 4-Supplement 1A), indicating that the trend was specific for TA cells. To assess whether the cell-type correlations were driven by any individual perturbation, perturbations were dropped one at a time from the dataset (dropping all single and pairwise conditions containing the perturbation) and correlations were re-calculated for each data subset. We found that dropping EGFR-i from the dataset largely abrogated the anticorrelation between TA number and EE fraction (Figure 4-Supplement 1B, red arrowhead), which is consistent with recent work suggesting that EE cell production can be triggered by EGFR-i-induced stem cell quiescence (Basak et al., 2017a). However, the anticorrelation between TA number and secretory fraction is not driven by a specific perturbation (Figure 4B), possibly indicating a tissue-intrinsic property.

**Figure 4.**
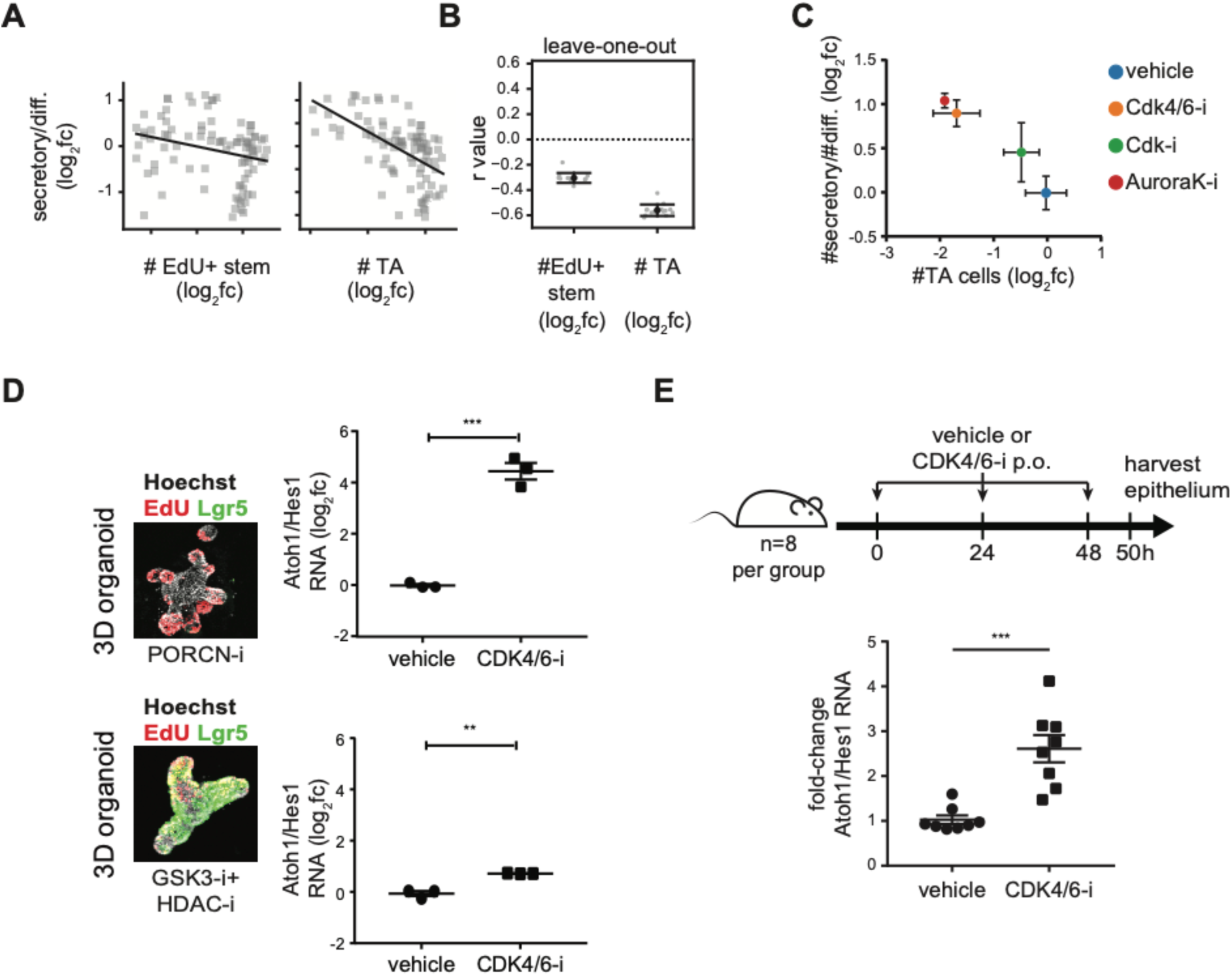
Inhibiting proliferation increases secretory cell prevalence in enteroid monolayers, in 3D organoids, and *in vivo*. (**A**) Numbers of transit-amplifying (TA) cells, but not EdU^+^ stem cells, correlate with secretory cell fractions. Perturbation effects (log_2_fc) are plotted pairwise for each feature. (**B**) The TA-secretory correlation is not driven by a specific perturbation. Each of 13 perturbations was sequentially dropped from the dataset and r value calculated. (**C**) Inhibiting cell cycle progression increases secretory cell fractions. Enteroids were treated as indicated for 48hrs, after which #TA cells, and #secretory/#differentiated cells were quantified. n = 3 wells. (**D**) TA cells are sufficient to alter secretory fractions in response to cell cycle inhibitors. 3D organoids were perturbed into stem or TA cell-rich states using PORCN-i (TA) or GSK3-i+HDAC-i (stem), respectively. These factors were removed and then a CDK4/6 inhibitor (palbociclib) was applied. The secretory:absorptive ratio was measured at 2 days post-treatment by qRT-PCR for the Atoh1/Hes1 ratio in TA-enriched (p=0.0002) or stem-enriched (p=0.01) 3D organoids. n = 3 wells. (**E**) Impairing proliferation increases the secretory:absorptive ratio *in vivo*. Mice were treated with CDK4/6-i (palbociclib) or vehicle every 24hr for 48hrs. At 50hrs, intestinal crypts were harvested and gene expression was measured by qRT-PCR. CDK4/6-i increases the Atoh1:Hes1 ratio (p=0.00005). n=8 mice/group. Error bars mean +/− sem.

We further investigated the anticorrelation between TA numbers and secretory cell fraction in enteroid monolayers, 3D organoids, and *in vivo*. To test whether the correlation is driven by TA proliferation, we directly modulated proliferation in enteroid monolayers using Cdk and Aurora kinase inhibitors and found that inhibiting proliferation, which decreases TA numbers, caused an increase in secretory cell fractions (Figure 4C). Further, CDK4/6 inhibition increased the secretory cell fraction (and decreased TA cell numbers) in a dose-dependent manner (Figure 4-Supplement 2). We then used 3D organoids to demonstrate that the correlation was specific to TA, and not stem cells. 3D organoid cultures were first enriched for either stem (treatment with GSK3-i+HDAC-i (Yin et al., 2014b)) or TA (treatment with PORCN-i) cells. qRT-PCR analysis confirmed enrichment for Lgr5 RNA over Ki67 RNA in GSK3-i+HDAC-i-treated 3D organoids, and Ki67 RNA over Lgr5 RNA in PORCN-i-treated 3D organoids (Figure 4-Supplement 3). IF staining also showed almost complete conversion to Lgr5^+^ stem cells in GSK3-i+HDAC-i-treated 3D organoids, while PORCN-i-treated 3D organoids retained proliferating (EdU^+^) cells but lost Lgr5^+^ stem cells (Figure 4D, images). Organoids were then removed from enrichment media and treated with the CDK4/6 inhibitor palbociclib, which significantly impairs proliferation in 3D intestinal organoids (Basak et al., 2017b). Atoh1 and Hes1 RNA levels were measured by qRT-PCR to quantify changes in secretory and absorptive cell fractions, respectively. Strikingly, CDK4/6 inhibition increased the ratio of Atoh1/Hes1 RNA in 3D organoids enriched for TA cells. However, CDK4/6 inhibition had a reduced effect on 3D organoids enriched for Lgr5^+^ stem cells (Figure 4D, graphs). These data indicate that proliferation of TA cells drives the observed changes in secretory prevalence. Lastly, we validated the correlation between TA proliferation and secretory fraction *in vivo*. Mice (n=8 per group) were treated with the CDK4/6 inhibitor palbociclib or vehicle. After 50 hours of treatment, intestinal crypts were harvested and gene expression was measured by qRT-PCR. CDK4/6-i treatment decreased Ki67 expression, indicating the treatment effectively decreased proliferation (Figure 4-Supplement 4A). Importantly, CDK4/6-i treatment increased the Atoh1/Hes1 ratio (Figure 4E) and, to a lesser extent, the Dll/Notch ratio (Figure 4-Supplement 4B), demonstrating that proliferation is anti-correlated with secretory fraction *in vivo*.

### Differential amplification as a model for proliferation-based control of tissue composition

How does altering TA cell proliferation affect the abundance of secretory cells relative to other differentiated (absorptive) cell-types? Previous studies suggest that secretory progenitors are less proliferative compared to absorptive progenitors. In particular, commitment to a secretory fate (specifically expression of the Notch ligand Dll1) has been observed to coincide with cell cycle exit (Stamataki et al., 2011), and lineage tracing experiments found that secretory cell clones are on average smaller than absorptive cell clones (Bjerknes & Cheng, 1999). To experimentally confirm that secretory progenitors indeed undergo fewer divisions than absorptive progenitors, an EdU dilution experiment was conducted. A pulse of EdU was administered for the first 9 hours of culture (less than one TA cell cycle length (Matsu-Ura et al., 2016)), followed by a chase of 39 hours (Figure 5A). EdU is initially incorporated into proliferating cells and then subsequently diluted with each cell division. Thus, EdU intensity in differentiated cells serves as a proxy for division number. Cells were fixed and stained for secretory cell markers Lyz, Muc2, and ChgA as well as EdU. EdU intensity distribution for all labeled cells was bimodal, indicating presence of two distinct subpopulations with one population of cells dividing fewer times (higher EdU intensity) than the other population (lower EdU intensity, Figure 5A). The EdU intensity distributions for Lyz^+^ Paneth, Muc2^+^ goblet, and ChgA^+^ EE cells were clearly shifted right (towards higher EdU intensity) compared to all epithelial cells, indicating that secretory cells divided fewer times compared to the overall population of intestinal epithelial cells (Figure 5A).

**Figure 5.**
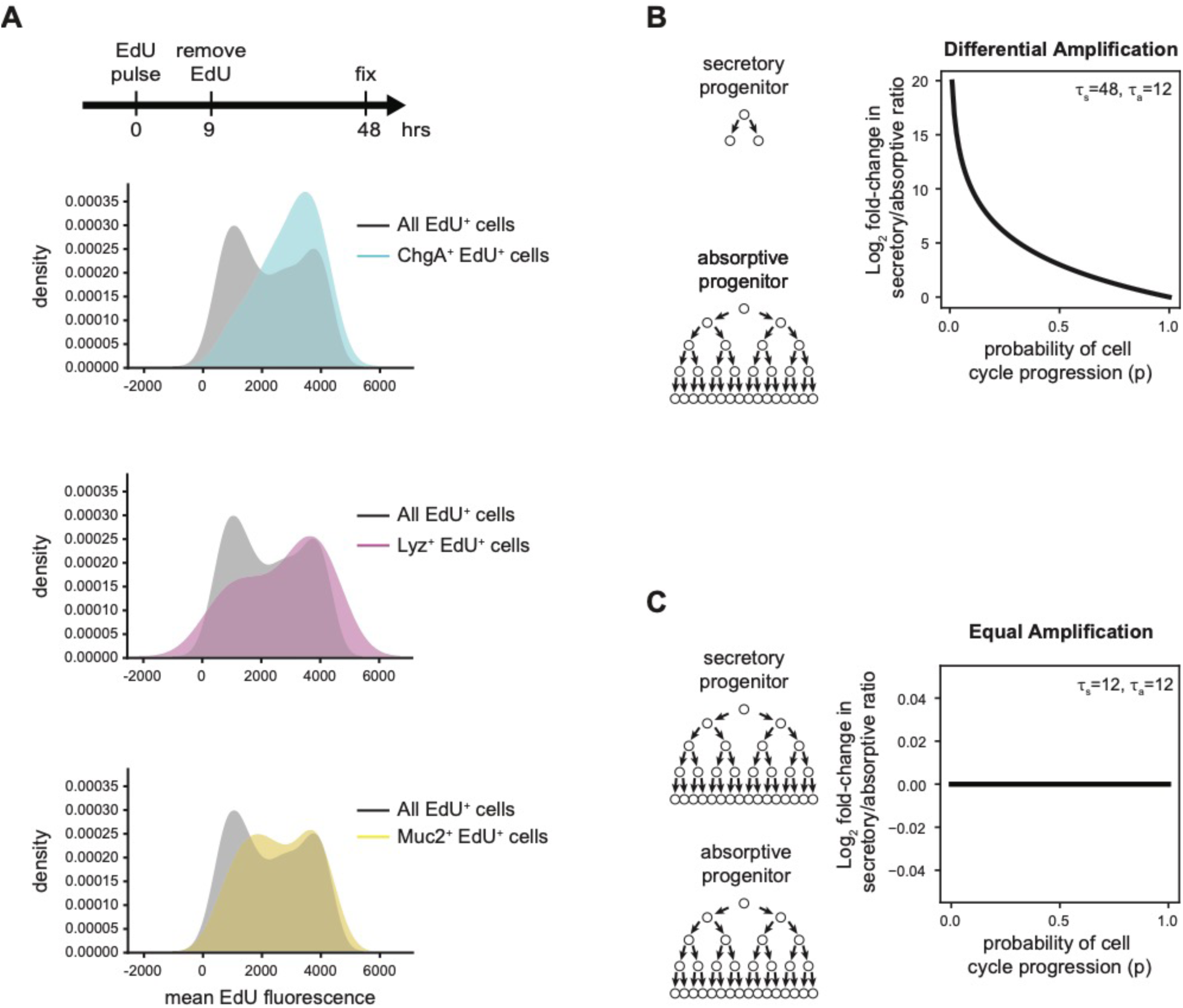
Differential amplification of secretory progenitors connects proliferation with differentiated cell-type composition. (**A**) Secretory progenitors divide fewer times than other progenitors. Enteroids were pulsed with EdU for 9 hours. After 39 further hours of culture enteroids were fixed and stained. Mean EdU signal intensity was quantified in all EdU^+^ cells and in EdU^+^ cells that also stained positive for markers of Paneth (Lyz), goblet (Muc2) and EE (ChgA) cells. Distribution of EdU intensities is represented as a kde plot. (**B**) If secretory progenitors divide fewer times than absorptive progenitors, inhibiting the cell cycle increases the secretory:absorptive ratio. Exponential growth model parameters are shown where secretory cell progenitors divide once (τ_s_=48hr) and absorptive cell progenitors divide four times (τ_a_=12hr) over a 48hr model time. (**C**) If secretory progenitors divide the same number of times as absorptive progenitors, inhibiting the cell cycle will not affect the secretory:absorptive ratio. Exponential growth model parameters are shown where both secretory and absorptive progenitors divide four times (τ_a_=τ_s_=12hr). For other model parameterizations see Figure 5-Supplement 1.

To mathematically evaluate whether differential amplification of secretory and absorptive progenitors could explain the observed anti-correlation between TA proliferation and secretory cell abundance, a simple exponential growth model was established. We computed the expected number of differentiated progeny cells from progenitor (TA) cells committed to either secretory fates or absorptive fates. The ratio of the expected output from secretory progenitors to that from absorptive progenitors captures the relative prevalence of secretory to absorptive cells in the differentiated population (Materials and Methods). To test how the relative difference in secretory and absorptive progenitor division numbers impacts the relationship between proliferation rate and tissue composition, we scanned through different division rates—and thus division numbers—for secretory and absorptive progenitors. We found that only parameter regimes where secretory progenitors divide fewer times than absorptive progenitors recapitulated the observed anti-correlation between proliferation and the ratio of secretory to absorptive cells (Figure 5B-C, Figure 5-Supplement 1). Thus, differential amplification of TA progenitors in the intestinal epithelium could provide a mechanism for controlling secretory-absorptive bias through proliferation.

## Discussion

How the intestinal epithelium globally integrates microenvironmental signals to regulate tissue composition has been challenging to study due to a lack of quantitative, high-throughput methods. Here, we developed a platform utilizing enteroid monolayers to comprehensively survey changes in intestinal epithelial cell-type composition to diverse perturbations. This enabled the identification of new culture conditions that enrich for specific cell-types, the dissection of perturbations that have unexpected combined effects on tissue growth, and the discovery that differential amplification of progenitor cells in different lineages can control differentiated cell-type composition. Importantly, recapitulation of perturbation effects and the proliferation-secretory link in both 3D organoids and *in vivo* lends support to the hypothesis generation ability of enteroid monolayers.

In terms of specific culture conditions, it was observed that pairwise combinations of GSK3, JAK1/2, HDAC, BMPR, and p38 MAPK inhibitors enriched for Lgr5^+^ stem cells. JAK1/2 inhibition has not been connected with regulation of stem cells under non-inflammatory conditions. Intriguingly, during inflammation JAK1/2 is essential for reserve stem cell activation (Richmond et al., 2018a). Since JAK1/2 inhibition increased Lgr5^+^ stem cell numbers, this could indicate a difference in the regulation of homeostatic (Lgr5^+^) and reserve stem cells. In addition, it was observed that combined TGF-beta stimulation and PORCN inhibition increases the fraction of enteroendocrine cells in the secretory lineage. TGF-beta signaling increases and Wnt signaling decreases as cells migrate up the crypt-villus axis (Barnard et al., 1989; Farin et al., 2016). Therefore, these data suggest a connection between spatial locations on the crypt-villus axis and differentiated cell-type composition.

By analyzing combinations of microenvironmental factors, we found more interactions amongst signals regulating progenitor (stem and TA) cell numbers than prevalence of differentiated cell-types. Possible reasons for the increased number of interactions include that either more signals control progenitor proliferation, or that signals act redundantly upon a smaller number of shared pathways. By examining specific pathway interactions, we found an unexpected mutual antagonism between IL-4 and EGFR-i that regulated the TA cell population. It was previously unknown that IL-4 induces BMP production, which has implications for its effects on the intestinal epithelium during type 2 immune responses such as worm infection (von Moltke et al., 2015). Further, it is surprising that IL-4 can bypass EGFR blockade, given recent data suggesting that EGFR is a gatekeeper of MEK-Erk-driven cell cycling in the intestinal epithelium (Basak et al., 2017a).

Lastly, our finding that perturbing TA cell proliferation altered the balance of secretory and absorptive differentiated cell-types underpins a new perspective on the role of the intestinal lineage structure in regulating cell-type composition of tissues. Lineage structures have been modeled in the past, such as the branching of progenitors in enabling robust feedback control (Lander et al., 2009). Here, we find that a differential amplification model of transit-amplifying progenitors in different lineages allows for control of tissue cell-type composition through modulation of proliferation. This model can be contrasted with a probabilistic stem cell decision model, in which cell-type composition is determined by stochastic decision-making in stem cells (Balázsi et al., 2011). We do not rule out that probabilistic fate decisions may still play an important role. However, the current study provides evidence that differential amplification of progenitors makes a significant contribution in shaping the tissue composition.

The link between TA proliferation and differentiated cell-type composition, as well as the observation of increased perturbation interactions in regulation of TA cells, point to a crucial and currently overlooked role for these short-lived progenitor cells in guiding tissue function. The intestinal epithelium, like many actively renewing tissues, is generated by stem cells that differentiate into short-lived progenitors, such as TA cells, and then assume differentiated cell fates. Intriguingly, many tissues, including skin and the hematopoietic system, feature transit-amplifying cells as intermediates between the stem and differentiated cell populations. By separately processing signals, these short-lived progenitor cell populations may broadly serve as a buffer that enables coordinated tissue responses to changing microenvironments (e.g. worm infections (Birchenough et al., 2016)) while insulating the stem cell population from extreme, transient changes.

## Materials and Methods

### Mice

All animal care and experimentation was conducted under protocol AN-179937 agreed upon by the Administrative Panel on Laboratory Animal Care at the University of California, San Francisco. All our animal studies are performed in full accordance with UCSF Institutional Animal Care and Use Committee (IACUC). 5- to 6-week-old male C57BL/6 mice (C57BL/6NHsd) were purchased from Harlan and housed with ad libitum food and water on a 12hr light cycle at the UCSF Preclinical Therapeutics Core vivarium.

### Media

Organoid basal media (OBM) consists of Advanced DMEM/F12 with non-essential amino acids and sodium pyruvate (Fisher Scientific #12634-028) containing 1x N-2 (Fisher Scientific #17502-048), 1x B-27 (Invitrogen #17504-044), 10 mM HEPES (Invitrogen #15630080), 1x GlutaMAX (Invitrogen #35050-061), 1 μM N-acetylcysteine (Sigma Aldrich #A9165), 100 U/mL penicillin and 100 μg/mL streptomycin (Corning #30-002).

For initial seeding, enteroid monolayers were maintained in OBM supplemented with 3 μM CHIR-99021 (Sigma Aldrich #SML1046), 50 ng/mL murine EGF (Invitrogen #PMG8043), 1 μM LDN-193189 (Sigma Aldrich #SML0559), 500 ng/mL murine R-spondin-1 (Peprotech #315-32), and 10 μM Y-27632 (Selleck Chemicals #S1049).

4 hours after initial seeding, media was changed into OBM supplemented with 50 ng/mL murine EGF, 100 ng/mL murine Noggin, and 500 ng/mL murine R-spondin-1. Perturbations applied in the studies described here were all applied in the background of this medium.

### Enteroid monolayer cultures

Enteroid monolayers were derived as previously described (Thorne et al., 2018). Briefly, jejunum was isolated from male mice between 6-12 weeks of age. Mice used were either from the C57BL/6 strain or, when indicated, the Lgr5^eGFP-DTR^ strain (H. Tian et al., 2011) (kind gift of Frederic de Sauvage, Genentech via Ophir Klein under MTA #OM-216813). Epithelium was released from jejunal tissue by incubation in ice-cold PBS with 3 mM EDTA in PBS (Ambion #9260). Released epithelial tissue was washed 3x with OBM, after which crypts were separated from villus material using 100 and 70 μm cell strainers (BD Falcon) in succession. Crypts were resuspended in seeding media and plated on Matrigel (Thermo Fisher #CB-40234C)-coated 96-well optical bottom plates (BD Biosciences #353219 and Greiner #655090). Typically, 300 crypts were seeded per well. We identified this seeding density because, at this density, we did not observe an effect of variations in initial confluency on cell outgrowth (#cells) or cell-type composition (Figure 1-Supplement 2). Four hours after seeding, cells were washed with OBM and incubated in control media containing other perturbations of interest.

### 3D organoid cultures

3D organoids were cultured as previously described (Sato et al., 2009). For imaging experiments, 3D organoids were seeded in 10 μL of Matrigel in 96-well optical bottom plates.

### CDK4/6-i administration to mice and tissue harvest

To test the effects of cell cycle inhibition on the secretory:absorptive ratio, palbociclib (LC Laboratories #P-7744) at 150 mg/kg in 50 mM sodium lactate buffer pH 4.4 was administered to mice by oral gavage every 24 hours for 48 hours (at 0 hours, 24 hours, and 48 hours). At 50 hours, the small intestine was harvested and intestinal crypts were harvested as described in the enteroid monolayer culture section above. Crypts were lysed in Buffer RLT (RNEasy Kit, Qiagen) for subsequent RNA purification.

### Growth factors and chemical compounds

All growth factors and chemical compounds were purchased from suppliers and used as designated without further purification. Unless otherwise indicated, perturbations were used as follows:

| **Perturbation** | **Vendor and Catalog #** | **Concentration** |
| --- | --- | --- |
| DAPT (Notch-i) | Stemgent #04-0041 | 10 μM |
| CHIR-99021 (GSK3-i) | Sigma Aldrich #1046 | 3 μM |
| Valproic acid (HDAC-i) | Sigma Aldrich #P4543 | 1 mM |
| IWP-2 (PORCN-i) | Selleck Chemicals #S7085 | 2 μM |
| Wnt3a | R&D Systems #5036-WN-500 | 200 ng/mL |
| TGF-β | PeptoTech #100-21C | 4 ng/mL |
| EW-7197 (TGF-βR-i) | Selleck Chemicals #S7530 | 1 μM |
| BMP4 | R&D Systems #314-BP-010 | 200 ng/mL |
| LDN-193189 (BMPR-i) | Sigma Aldrich #SML0559 | 1 μM |
| Baricitinib (JAK1/2-i) | Selleck Chemicals #S2851 | 2 μM |
| SB202190 (p38MAPK-i) | Sigma Aldrich #S7067 | 10 μM |
| IL-4 | PeptoTech #214-14 | 20 ng/mL |
| Erlotinib HCl (EGFR-i) | Selleck Chemicals #S1023 | 2.5 μM |
| PD0325901 (MEK-i) | Selleck Chemicals #S1036 | 1 μM |
| Palbociclib (PD-0332991) (CDK4/6-i) | Selleck Chemicals #S1116<br>(organoid studies) or<br>LC Laboratories #-7744<br>(*in vivo* studies) | 10 μM |
| Flavopiridol (CDK-i) | Selleck Chemicals #S2679 | 0.3 μM |
| AT9283 (AuroraK-i) | Selleck Chemicals #S1134 | 1 μM |

### Immunofluorescence assay

#### Enteroid monolayers

Enteroid monolayers were washed 1x with warm D-PBS and then fixed with 4% paraformaldehyde in PBS for 15min at room temperature. Cells were then washed with PBS and permeabilized with 0.5% Triton-X-100 in PBS at room temperature for 10min. Cells were washed, blocked with 3% BSA in PBS for 30min, and then incubated in primary antibody in antibody buffer (PBS with 0.3% Triton-X-100, 1% BSA) overnight at 4C. The next day, cells were washed and incubated with secondary antibodies and Hoechst 33342 (5 μg/mL; Invitrogen #H3570) in antibody buffer for 2 hours at room temperature. After this, cells were washed with PBS and imaged in TBS-T (0.1% Tween in 1x TBS pH 7.4).

#### 3D organoids

Media was carefully aspirated from around Matrigel domes containing 3D organoids using a P100 pipet. 4% paraformaldehyde in PBS was immediately added for 15min at room temperature. Cells were then washed 2x with PBS and permeabilized using 0.5% Triton-X-100 in PBS for 20min at room temperature. Cells were then rinsed 3×10min with 100 mM glycine in PBS with gentle agitation. Cells were blocked in 3% BSA in PBS for 40min and then incubated with primary antibody in antibody buffer overnight at room temperature. The next day, cells were washed 3×20min in antibody buffer and then incubated with fluorescent secondary antibodies and Hoechst in antibody solution for 1hr at room temperature. Cells were then rinsed in PBS and stored and imaged in TBS-T.

### Antibodies

All antibodies were purchased from suppliers and used as designated without further purification. Unless otherwise indicated, antibodies were used as follows:

| Epitope | Vendor and Catalog # | Dilution |
| --- | --- | --- |
| Lysozyme (Lyz) | Dako | 1:2000 |
| Mucin-2 (Muc2) | Santa Cruz Biotechnology #15334 | 1:100 |
| Chromogranin A (ChgA) | Santa Cruz Biotechnology #393941 | 1:100 |
| GFP | Abcam #5450 | 1:2000 |
| Dclk1 | Abcam #31704 | 1:1000 |
| Erk1 (pT202/pY204) + Erk2 (pT185/pY187) (pErk) | Abcam #50011 | 1:200 |
| Cyclin D1 | Life Technologies #MA5-14512 | 1:200 |

### EdU pulse and visualization

To visualize proliferating cells (specifically, those in S phase), enteroid monolayers were incubated with 10 μM EdU (Thermo Fisher #A10044) in media (containing indicated perturbations or vehicle) for 2 hrs prior to fixation. After immunofluorescence staining, EdU^+^ cells were visualized using Click chemistry as previously described (Salic & Mitchison, 2008). Briefly, cells were incubated with a reaction mixture containing 1 mM CuSO4 (VWR International #470300-880), 5 μM sulfo-Cyanine5 azide (Lumiprobe #B3330) or 5 μM BDP-FL azide (Lumiprobe #11430), and 100 mM sodium ascorbate (Sigma Aldrich #A4034) in PBS for 30min at room temperature.

### Automated brightfield microscopy

Upon initial plating, enteroid monolayers were imaged in the brightfield channel using the 10x objective of a Nikon TE200-E epifluorescence microscope. These data were used as a control to determine whether enteroid monolayers were seeded at an optimal and consistent confluency.

### Quantifying % confluency

% Confluency (percent of image which is occupied by enteroid monolayer cultures) was quantified from brightfield images using a previously reported algorithm (CellularRegionsFromBrightField function in Supplementary Software 1 from reference (Ramirez et al., 2016)).

### Automated confocal microscopy

Enteroid monolayers were imaged on the 10x objective of a Nikon A1 confocal with Ti2-E microscope. The area of each well was covered by 24 individual scans. In each field of view, 4-8 z planes were collected at 1024×1024 resolution. Importantly, the nuclear stain was used to autofocus in each new field of view.

### Data availability

All data generated or analyzed during this study are included in this published article (and its supplementary information files)

### Immunofluorescence image segmentation and analysis

#### General information

Image segmentation was performed using a custom Python analysis pipeline. All code and documentation can be found on GitHub (https://github.com/AltschulerWu-Lab/MAGS). Starting with maximum intensity projections of Hoechst, EdU, Lgr5-GFP, Muc2, Lyz, and ChgA fluorescent images, we segmented and then quantified numbers of nuclei, EdU^+^ cells, stem cells, goblet cells, Paneth cells, and enteroendocrine cells, respectively. The general segmentation process for each object type consisted of two major steps: a thresholding step to identify image foreground, and a segmentation step to generate location and boundary of objects. Specific details are as follows:

#### Segmenting nuclei

Hoechst stain images were smoothed through convolution with a bilateral filter. The foreground was identified using a modified Otsu threshold method. Nuclei in enteroid cultures are highly heterogeneous with small, tightly packed nuclei in the crypt regions and large, sparsely distributed nuclei in the villus regions. Therefore, to obtain an accurate segmentation in these different regions, we conducted a two-step segmentation approach. First, sparse nuclei were identified using a multi-scale Laplacian of Gaussian (LoG) detector to generate markers of object locations. A watershed algorithm was then used to generate object boundaries. Clumps were detected in the sparse segmentation using object size and shape irregularity cutoffs. These clumps contained mis-segmented dense nuclei and were subsequently segmented using differently parameterized LoG detector to identify object seeds and watershed to generate object boundaries. The dense and sparse segmentations were then merged for the final result (also see Figure 1-Supplement 1C).

#### Segmenting EdU^+^ nuclei

The same process as nuclear segmentation was used because nuclear objects in EdU stain images had similar properties as those in Hoechst stain images.

#### Segmenting stem cells (Figure 1-Supplement 1D)

GFP immunofluorescence in enteroid monolayers derived from Lgr5-eGFP-DTR mice (H. Tian et al., 2011) revealed both crypt regions, with multi-cell membrane GFP staining, and single GFP^+^ cells outside crypt regions. The single GFP^+^ cells appeared morphologically similar to tuft cells and co-stained for Dclk-1 (Figure 1-Supplement 1E), indicating that these cells are likely a subpopulation of tuft cells rather than stem cells. Therefore, we only identified stem cells within crypt regions. Crypt regions were defined as clusters of Lgr5^+^ stem and Paneth cells, corresponding to crypt bases *in vivo*. Lgr5-GFP stain images were processed to remove tissue background and thresholded to identify foreground. Holes and gaps in crypt regions were filled using morphological operations and small objects (typically, Lgr5^+^/Dclk1^+^ cells, see Figure 1-Supplement 1E) were dropped. The remaining objects were labeled as crypts. We then counted nuclei within crypt regions as stem cells. Importantly, we removed any nucleus in a crypt region that was associated with a Paneth cell from the stem cell count (Figure 1-Supplement 1A).

#### Identifying Paneth cells

Lysozyme (Lyz) immunofluorescence images were smoothed through convolution with a bilateral filter then a tophat filter. Foreground was identified using the Otsu-thresholded Lyz immunofluorescence image. A LoG detector was then used to generate markers of Paneth object locations.

#### Segmenting goblet cells

Mucin-2 (Muc2) stain images were smoothed by convolution with a median filter. Foreground was identified using a convex hull of objects in each Otsu-thresholded Muc2 immunofluorescence image. Goblet cells were segmented using a LoG detector to generate markers of goblet object locations followed by watershed to create object boundaries.

#### Identifying enteroendocrine (EE) cells

ChgA stain images were processed using the same steps as Paneth cell identification, only with different parameters.

#### Evaluation of Image Segmentation

Each cell-type object (e.g., each nucleus, each goblet cell, each Paneth cell) was identified in raw immunofluorescence images by hand by an expert and, in parallel, using the customized algorithms described above. The expert-generated segmented images (where each mask represents an individual object) were compared to algorithm-generated segmented images to determine algorithm performance. ‘Precision’ was quantified by dividing the number of true positives (expert-identified objects also identified by the algorithm) by the number of total positives (all algorithm-identified objects). ‘Recall’ was quantified by dividing the number of true positives by the total number of expert-identified objects. F1 scores were calculated as the harmonic mean of precision and recall. See Table 1 for results.

### Data Analysis

#### Extracting numbers of each cell-type

Due to limitations of conventional fluorescence microscopy, all desired cell-types were quantified across two replicate immunofluorescent stain sets: a crypt stain set (Hoechst, EdU, Lgr5-GFP, Lyz) and a differentiated cell stain set (Hoechst, EdU, Muc2, ChgA). We were able to quantify the numbers of some cell-types directly from a single stain set, including the number of each directly measured cell-type (EdU^+^, stem, goblet, Paneth, EE) as well the number of proliferating (EdU^+^) stem cells (combining information from EdU and Lgr5-GFP segmentation), transit-amplifying cells (TA; #EdU^+^ cells minus #stem cells), and differentiated cells (# cells minus stem and TA cells).

#### Replicates and Error Estimation

In-plate replicate control wells (2-6 wells per plate) were used to estimate mean and error. For replicate plates, mean and error were pooled. For across stain-set readouts, error was propagated.

#### Fold Change

We calculated fold-change effects relative to in-plate controls for readouts within each stain set (#EdU^+^ stem cell, #TA cells). Fold-changes for readouts calculated across both stain sets (#secretory/#diff, #goblet/#secretory, #Paneth/#secretory, #EE/#secretory) were calculated to a pooled control baseline measurement.

### Statistical testing

In comparing enteroid samples, two-sided Welch’s t-tests were performed and p values were reported. In comparing *in vivo* samples, two-sided Student’s t tests were performed and p values were reported. Degrees of freedom is taken as n-1. Measurements were taken from distinct samples.

### Perturbation effect visualization

For Figure 2B, double perturbation phenotypes were sorted into similar phenotypes using hierarchical clustering (clustermap function in seaborn) with a euclidean distance metric. Single perturbation phenotypes were sorted based on the number of EdU^+^ stem cells in each row.

### Population growth model

We established a simple population growth model to compute the changes in secretory and absorptive populations in response to cell cycle perturbation. In this model, we assumed that the initial ratio of absorptive to secretory TA progenitors produced by stem cells is equal. We further assumed that the progenitors are locked into either secretory or absorptive fates after the initial commitment with no switching occurring between fates (Stamataki et al., 2011; van Es et al., 2012). The model describes the theoretical output of differentiated cells from populations of initial secretory or absorptive progenitors.

Under normal growth (control condition), we describe the amplification of absorptive and secretory TA cells as follows:

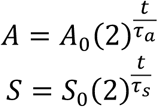

where A_0_ and S_0_ are the numbers of initial absorptive and secretory TA progenitors, respectively. A and S are the numbers of final absorptive and secretory differentiated cells, respectively. t is the total model time, set to be 48 (hours). τ_a_ and τ_s_ are the cell cycle lengths of absorptive and secretory progenitors, respectively. Absorptive progenitors are generally found to divide 4-5 times while secretory progenitors only divide 1-2 times (Potten, 1998; Stamataki et al., 2011; van Es et al., 2012). We set τ_a_ and τ_s_ to be 12 and 48 (hours) in control conditions. Thus, secretory progenitors divide once while absorptive progenitors divide four times before fully differentiating, consistent with previous observations. Progenitors that divide fewer times are not considered differentiated cells.

We model the perturbation effect with the parameter p, where p is the probability of cell cycle progression for any TA cell (alternatively 1 – p is the probability of quiescence). The expected numbers of differentiated absorptive and secretory cells, as averaged over an ensemble of initial progenitor populations, are as follows:

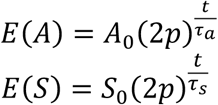

where 0 < p < 1 in the case of cell cycle inhibitors and p = 1 in the untreated case. Scanning through different values of τ_a_ and τ_s_, we observe that as long as τ_a_ is less than τ_s_, decreasing proliferation (increasing p) corresponds to an increase in secretory prevalence. The expectation follows from the observation that *p* modifies the expected number of progeny per generation to be *2p*. More explicitly, the expectation of progeny number (*X_i_*) for progenitor generation *i* can be calculated as follows:

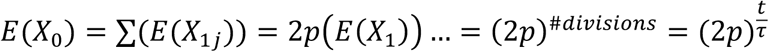

We note that we can take a different parametrization for the cell-cycle inhibitor effect where the length of cell cycling is varied under perturbation. The equations for this parametrization are as follows:

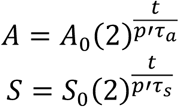

where we take p’ to be a number > 1 in the case of cell cycle inhibitors and p = 1 in the untreated case and t is 48 as before. In this approach, we do not lose progenitors to quiescence and all progenitors are considered differentiated at the end of the experiment time t. Since we can convert between p’ and p by letting 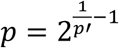, we would observe a similar outcome as the first parametrization approach where increasing perturbation effect via increased cell cycle inhibition corresponds to an increase in secretory prevalence.

### Identifying perturbation interactions

#### Multiplicative model

To identify perturbation interactions, we first established a model for how perturbation effects should combine if there is no interaction. A simple model commonly used in genetic and drug interaction studies is the multiplicative model (Mani et al., 2008; van Hasselt & Iyengar, 2019). Under the multiplicative model, perturbations that do not interact should combine as the product of their respective fold-change effects. This is equivalent to log-additivity, which can be calculated as follows:

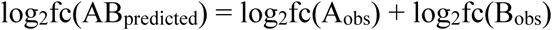

Features with 0 values had a pseudocount of 1 instead. Perturbation combination effects that deviate from the multiplicative model indicate a potential interaction, which we define biologically as the due to utilization of a shared pathway or redundant pathways (Dixon et al., 2009).

For each perturbation pair, we evaluated how a multiplicative model fits the observed double perturbation effects for each cell-type readout (e.g., #TA cells). To evaluate the multiplicative model across each perturbation and readout, we plotted the effects of perturbation combinations (e.g, the effects of Wnt3a in combination with all other perturbations on #TA cells) against the single perturbations (e.g., all perturbations except Wnt3a) in log scale. The multiplicative model prediction is a line offset on the y-axis by the effect of the perturbation of interest (Figure 3-Supplement 1B). Combinations of perturbations that do not exhibit an interaction will have an effect that lies on this line. We compared linear and multiplicative functions for each single perturbation and observed that some single perturbations fit well with either multiplicative or linear fit models, while others deviated significantly from both (e.g., Wnt3a fits well with either model while EGFR-i deviates significantly from both, see Figure 3-Supplement 1B-C). We computed the residuals for perturbation combination fit across all readouts. The two models had similar distributions of residual values with peaks close to zero (Figure 3-Supplement 1A), indicating that the majority of perturbations in our experiment combined in a manner that is well described by the multiplicative model.

#### Effect size

The deviation of each combinatorial perturbation from the prediction of the multiplicative model is represented as effect size, which was quantified in this case as absolute Cohen’s d, or the standardized deviation of the combinatorial phenotype from multiplicative model prediction.

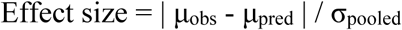

where μ_obs_ and μ_pred_ are the observed and predicted means for the combination effect and σ_pooled_ is the pooled standard deviation. In Figure 3A, the cut-off for the effect size was 5.

### BMP2 ELISA

Supernatant levels of BMP2 were quantified using a BMP-2 Quantikine ELISA kit (R&D Systems #DBP200), without significant deviations from manufacturer’s instructions.

### qRT-PCR

RNA was harvested from enteroid monolayers using an RNEasy Plus Mini Kit (Qiagen #74136). Reverse transcription was performed using iScript Reverse Transcription kit (Bio-Rad #1708841). Quantitative PCR was performed using SsoAdvanced Universal SYBR Green Supermix (Bio-Rad #1725272) on a BioRad CFXConnect. Test gene values were normalized to beta-actin values. To quantify Atoh1/Hes1 ratio, both Atoh1 and Hes1 fold-changes relative to control were calculated and then Atoh1 fold-change was divided by Hes1 fold-change. RNA levels were determined using the following primers:

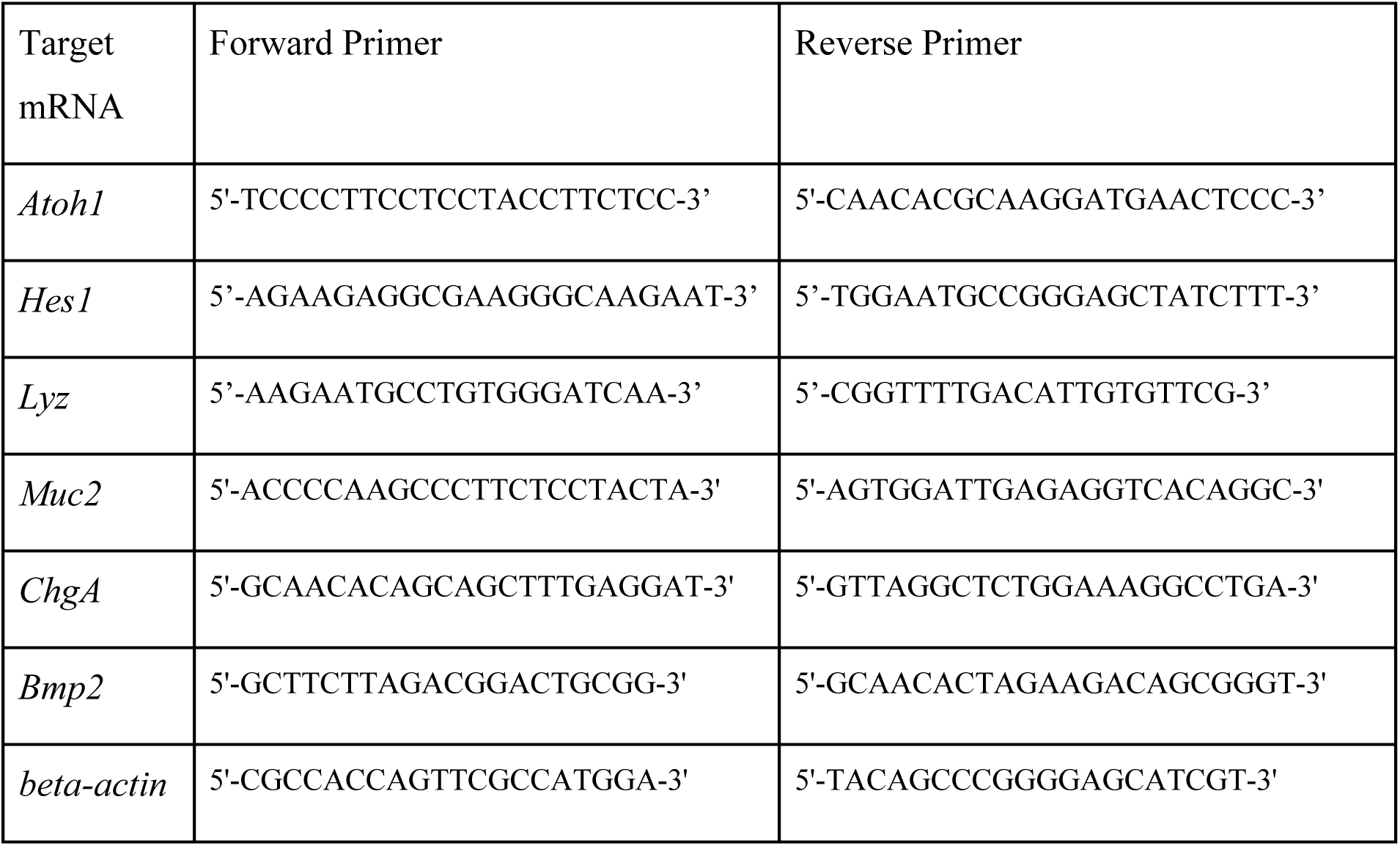

## Acknowledgments

We are grateful to Zev Gartner, Ophir Klein, Curtis Thorne, and members of the Altschuler-Wu lab for their feedback. We additionally thank Ophir Klein and Frederic de Sauvage for mouse resources. This work was supported by NIH GM112690 (S.J.A.), NCI-NIH RO1 CA184984 (L.F.W.), the UCSF Program for Breakthrough Biomedical Research which is partly funded by the Sandler Foundation (L.F.W.), NIH NRSA fellowship F32DK120102 (L.E.S.), and NSF GRFP fellowship 1650113 (I.W.C.).

## Author contributions

I.W.C. and L.E.S. conceptualized study, developed computational methods, designed and executed experiments, analyzed data, and wrote the manuscript. J.M.B. and V.S. designed and executed *in vivo* experiments and analyzed data. B.H. supervised the *in vivo* study. L.F.W. and S.J.A. conceptualized and supervised the overall study and wrote the manuscript.

---

**Figure 1-Supplement 1.**
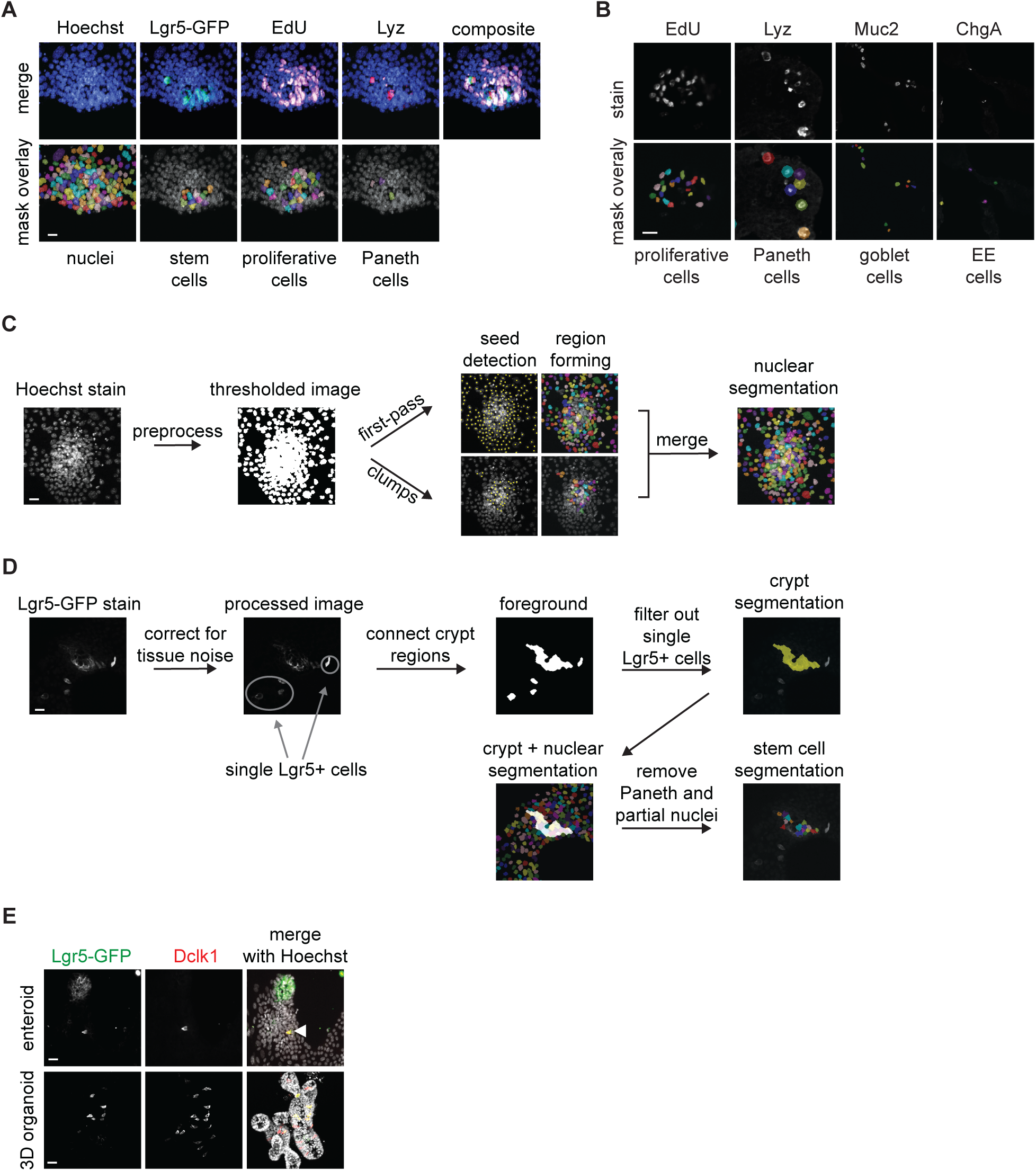
Quantification of intestinal epithelial cell-types in immunofluorescence images of enteroid monolayers. (**A**) Example of crypt segmentation with pseudocolor overlay for each cell-type readout. Scale bar 10μm. (**B**) Example of segmentation of proliferating, Paneth, goblet, and enteroendocrine (EE) cells. Top is raw immunofluorescence image for indicated marker and bottom includes pseudocolor overlay of each identified cell object. Scale bar 10μm. (**C**) Schematic of nuclear segmentation steps. Thresholded Hoechst stain images were segmented in two passes. The first pass segmented sparse nuclei and the second pass segmented clumped nuclei. Sparse and clumped segmentation were merged into the final nuclear segmentation. Yellow dots indicate identified markers of nuclear object locations, multi-pseudocolor overlay depicts individual nuclei segmented using a watershed algorithm. Scale bar 10μm. (**D**) Schematic of stem cell segmentation. The Lgr5-GFP stain was first corrected for tissue noise and then thresholded. Size filtering was used to separate multi-cell membrane GFP regions (crypt regions) from single Lgr5-GFP+ cells. All nuclei in crypt regions (nuclei identified from Hoechst image segmentation of same region), with the exception of Paneth cell-associated nuclei, were counted as stem cells. Scale bar 10μm. (**E**) Single Lgr5-GFP+ cells are also Dclk1+ in enteroid monolayers (top; scale bar 10μm) and 3D organoids (bottom; scale bar 15μm) and are thus excluded from stem cell counts.

**Figure 1-Supplement 2.**
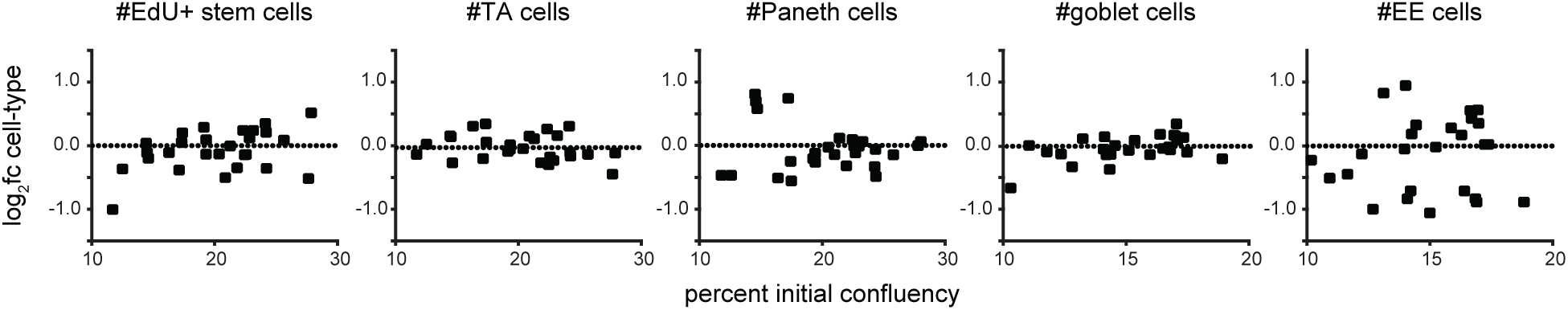
Enteroid monolayers exhibit reproducible cell-type composition across replicate wells when crypts are seeded at 10-30% initial confluency. Percent initial confluency was measured from brightfield images taken after crypt seeding and 48hr cell-type composition was quantified from immunofluorescence images. Y axis log_2_fc was computed relative to the average of all wells. No relationship is observed between initial confluency and cell frequencies.

**Figure 2-Supplement 1.**
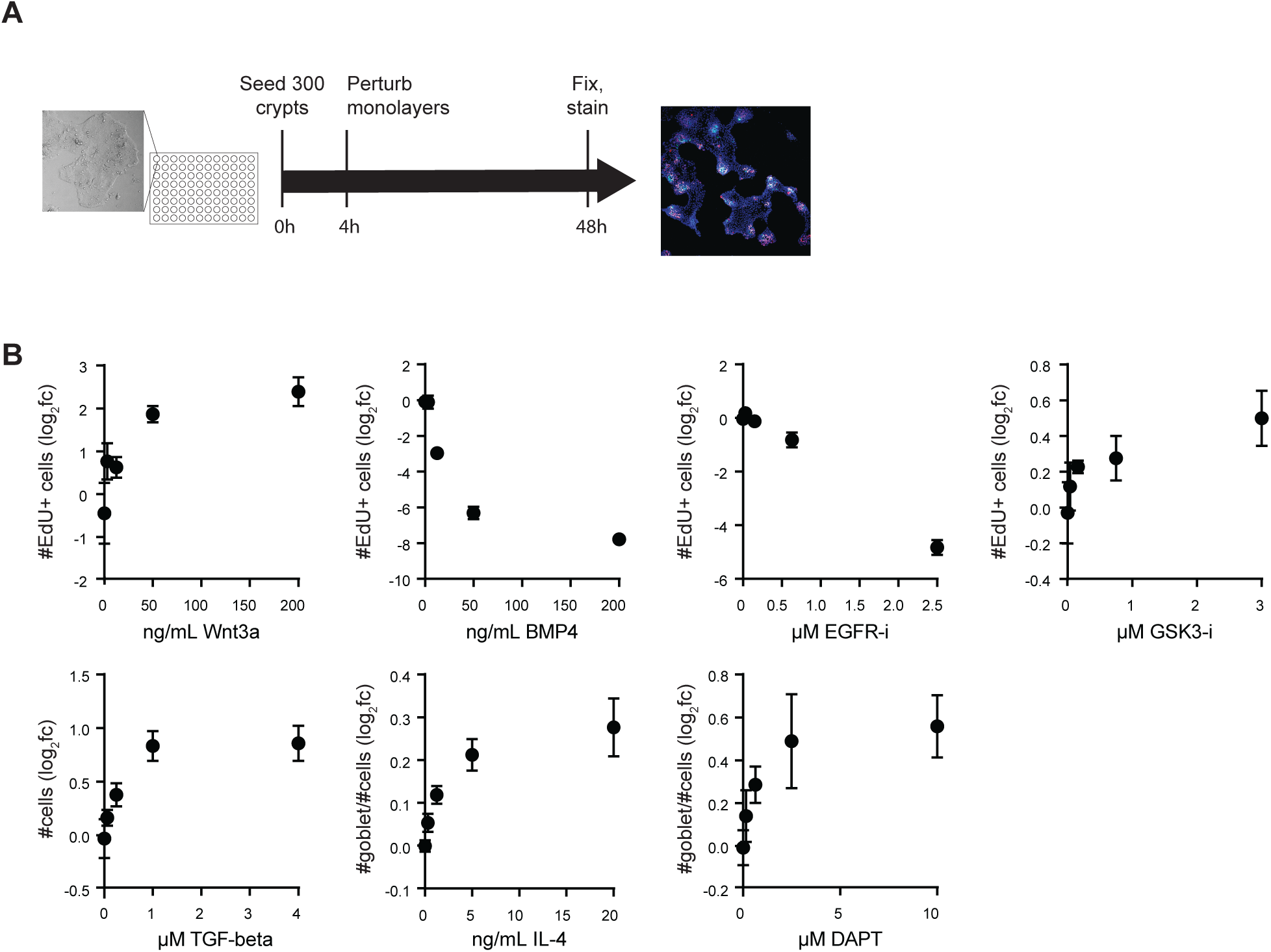
Optimization of perturbation dose. (**A**) Schematic of experimental procedure for investigating tissue responses to perturbation. (**B**) Quantification of cell numbers and prevalence after perturbation with increasing doses of Wnt3a, BMP4, EGFR inhibitor, GSK3 inhibitor, TGF-beta, IL-4, or Notch inhibitor. All data are represented as log_2_ transform of the fold-change effect relative to control. Fraction of goblet cells was calculated as a percentage of all cells for comparison with previous studies. n=2-6 wells. Error bars mean +/− sem.

**Figure 2-Supplement 2.**
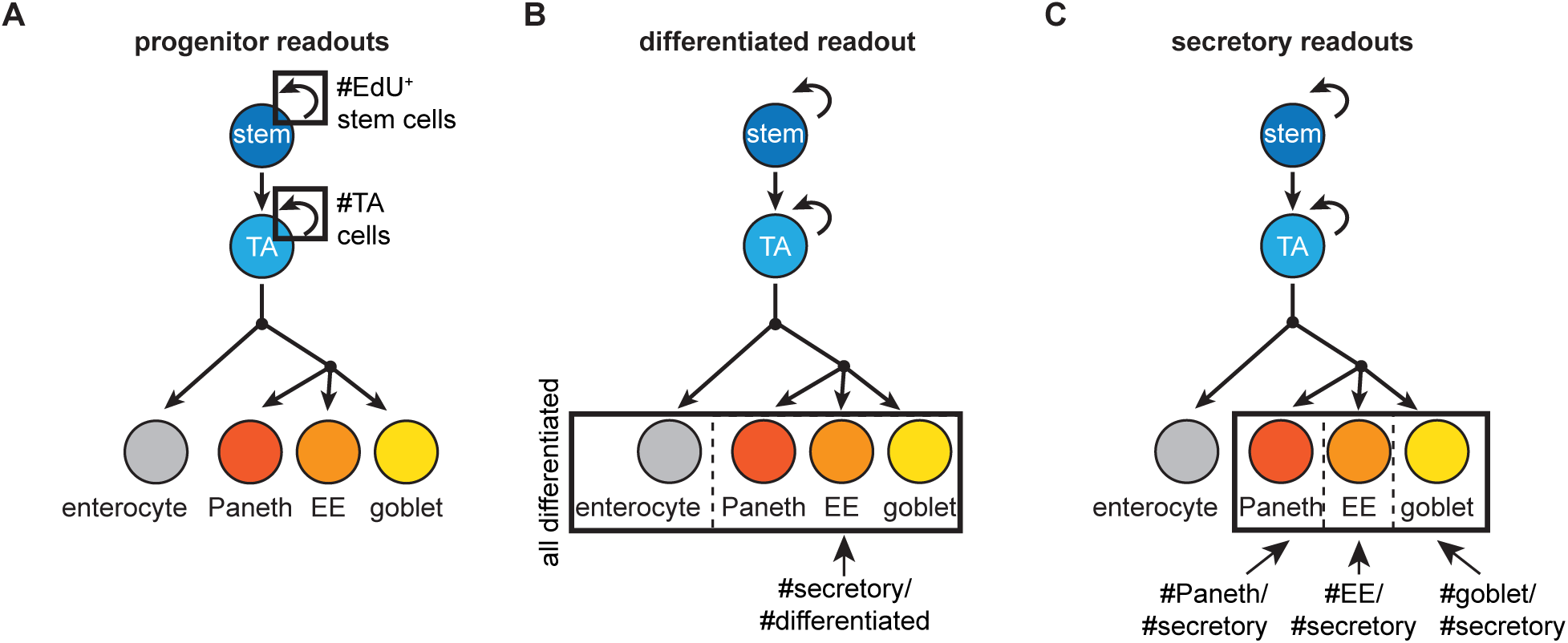
Combining cell-type numbers into readouts of progenitor proliferation and differentiated cell composition. (**A**) Numbers of proliferating progenitors were quantified as the #EdU^+^ cells, which was further divided into proliferating stem and transit-amplifying (TA) cells based on Lgr5 staining. (**B**) The secretory-absorptive bias was quantified as the fraction of differentiated cells (EdU^-^ and Lgr5^-^) that express secretory markers (the total number of cells that express Paneth (Lyz), goblet (Muc2), or enteroendocrine (EE) cell (ChgA) markers). (**C**) Composition of the secretory lineage was quantified as the fraction of secretory cells that express either goblet, Paneth, or EE cell markers.

**Figure 2-Supplement 3.**
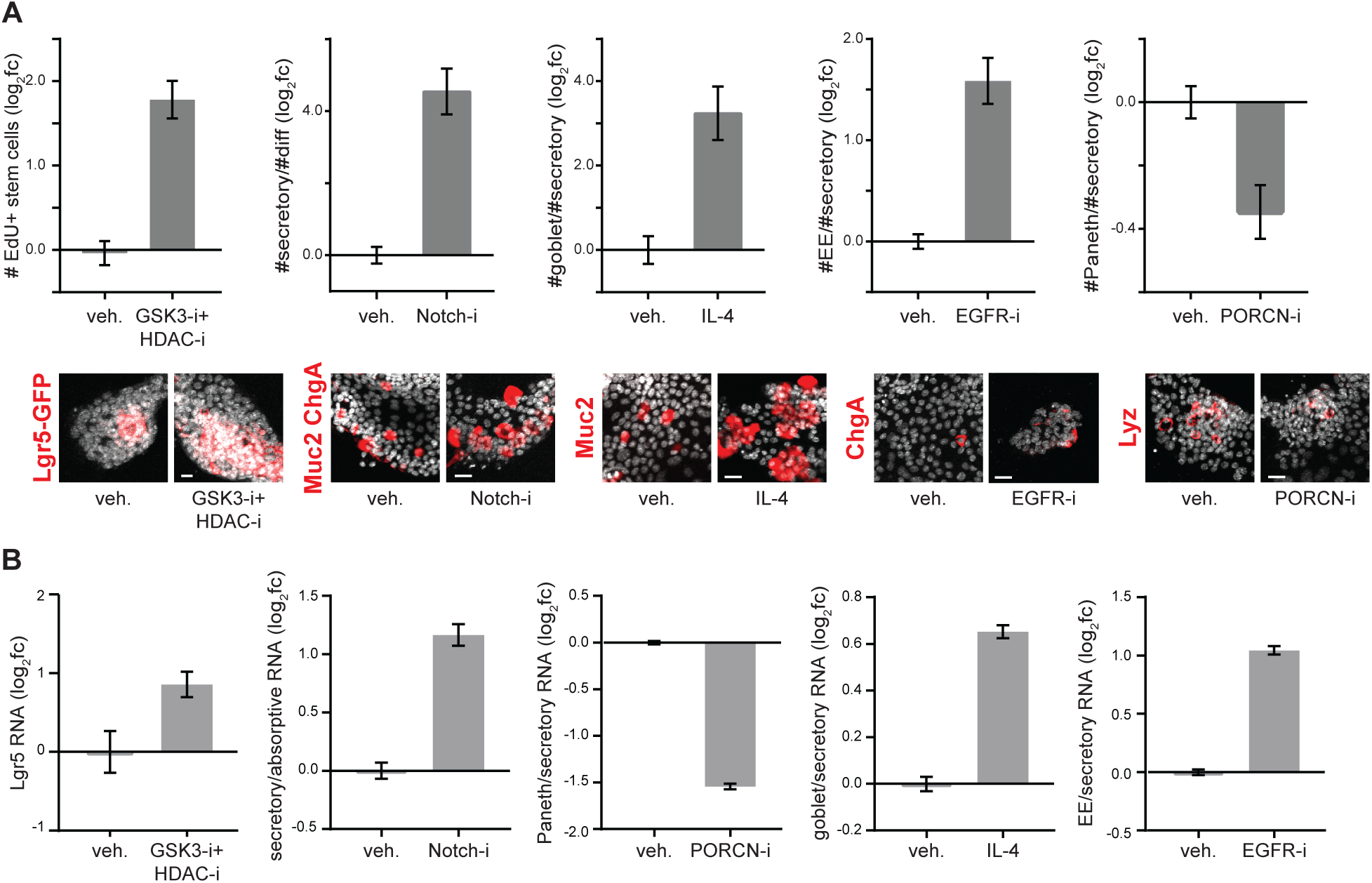
Benchmarking enteroid monolayer responses to well-characterized perturbations. (**A**) Enteroid monolayers respond as expected to modulators of stemness (GSK3-i+HDAC-i) and secretory cell prevalence (Notch-i, IL-4, EGFR-i, PORCN-i). Both quantification of replicate wells (top, n=2-6 wells) and representative images (bottom) are shown. Scale bars 10μm. (**B**) Similar changes in cell-type composition are observed at the RNA level. Enteroid monolayers were treated as indicated for 48 hours. For secretory:absorptive RNA, the Atoh1/Hes1 ratio was measured. Paneth RNA=Lyz, goblet RNA=Muc2, EE RNA=ChgA, secretory RNA=Lyz+Muc2+ChgA. n=3 wells. Error bars mean +/− sem. Veh=vehicle.

**Figure 3-Supplement 1.**
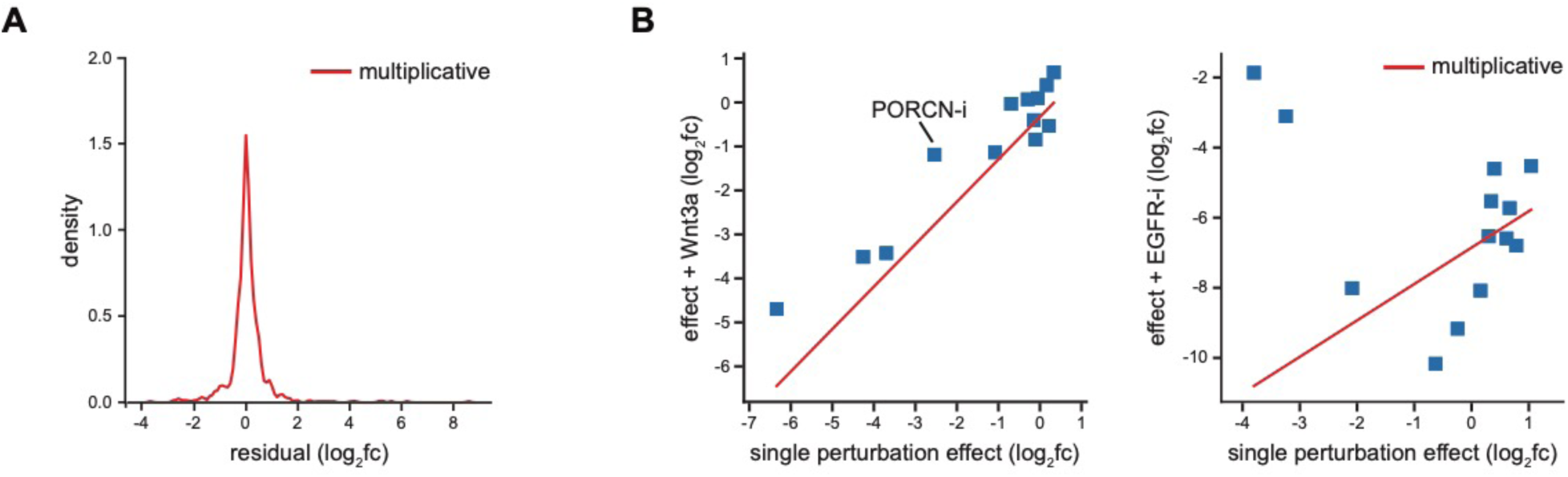
Evaluation of the multiplicative model. (**A**) Perturbation effects are generally similar to the predictions of the multiplicative model. Distribution of residual between predicted and observed values across all cell-type readouts is shown. (**B**) Individual perturbations are either predicted by the multiplicative model or diverge significantly. Examples of well-predicted (Wnt3a) and divergent (EGFR-i) perturbations are shown. Effects shown are on #TA cells. The multiplicative model predicts an interaction between an inhibitor of PORCN, an enzyme required to process epithelial-intrinsic Wnt3a, and Wnt3a treatment (left).

**Figure 3-Supplement 2.**
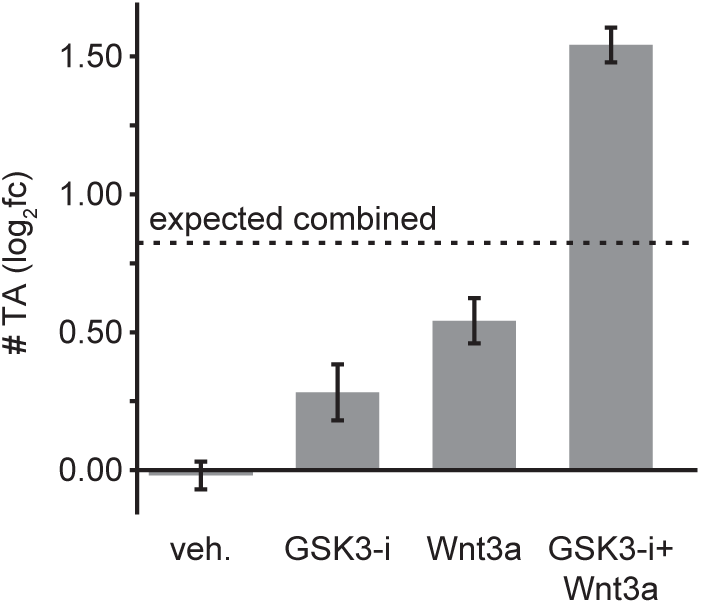
Example of perturbation interaction between Wnt3a and GSK3-i. The combination effect of Wnt3a and GSK3-i is much higher than would be expected under the multiplicative model. This synergism can be explained by the mechanisms of Wnt3a and GSK3-i as both perturbations impact components of the Wnt pathway.

**Figure 3-Supplement 3.**
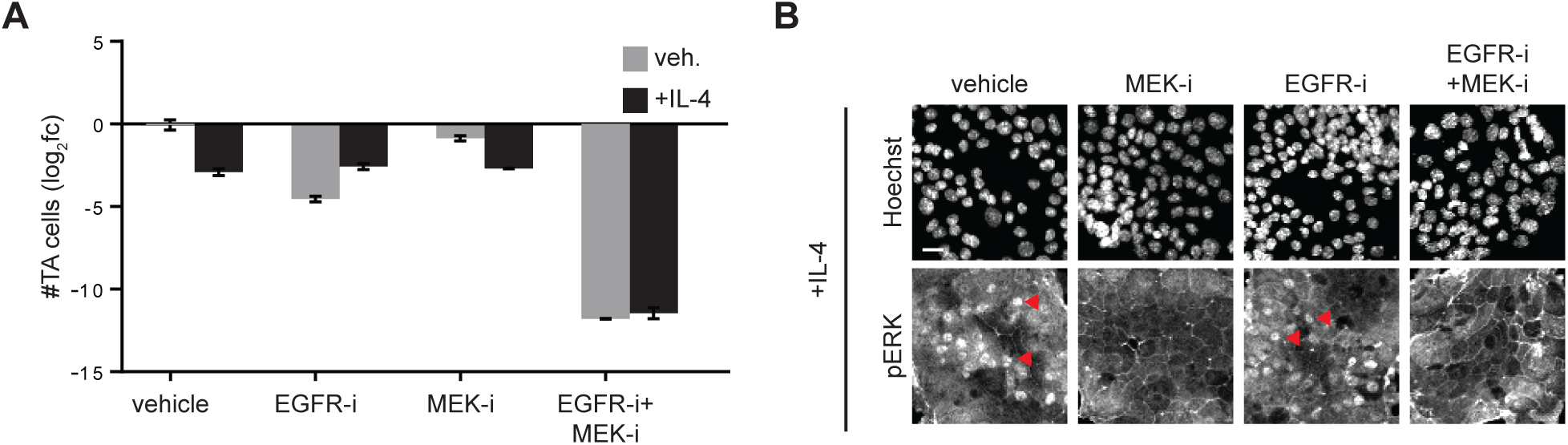
Regulation of MEK-Erk activity by EGFR and IL-4. (**A**) Antagonistic effect of IL-4 on EGFR-i is dependent on MEK activity. Enteroid monolayers were treated as indicated for 48 hours and the number of TA cells quantified. n=2-6 wells. (**B**) MEK inhibition reduces Erk activation (nuclear translocation of phospho-Erk) in enteroid monolayers even in the context of IL-4 treatment. Enteroid monolayers were treated with the indicated compounds for 24hr and then stained for phospho-Erk1/2. Red arrowheads indicate example cells with nuclear phospho-Erk. Scale bar 7.5μm.

**Figure 3-Supplement 4.**
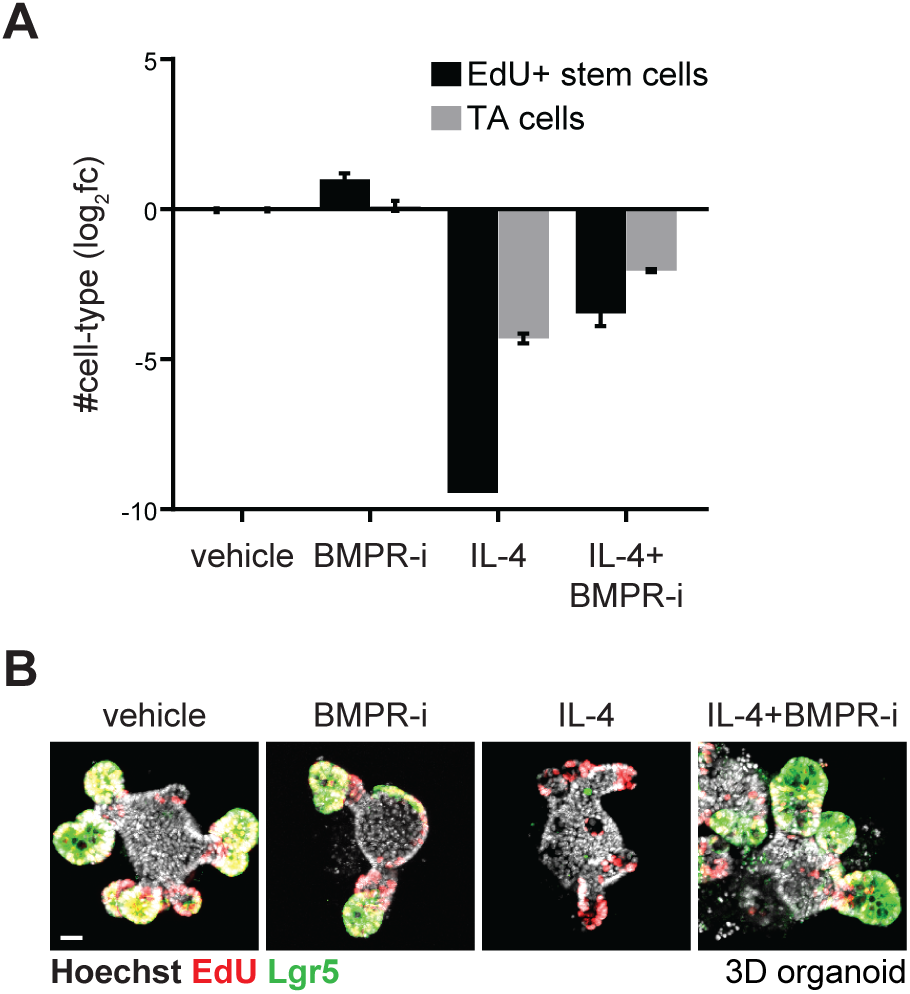
Regulation of BMP signaling by IL-4. (**A**) BMP receptor inhibition blocks IL-4-induced downregulation of TA cell numbers and stem cell numbers. Enteroid monolayers were treated as indicated for 48 hours and both EdU^+^ stem and TA cell numbers quantified. n=2-6 wells. (**B**) BMP receptor inhibition blocks IL-4-induced downregulation of proliferation and stemness in 3D organoids. 3D organoids were treated with vehicle or IL-4 in the presence and absence of BMP receptor inhibitor for 48hrs and then stained for proliferating cells (EdU^+^) and stem cells (Lgr5^+^). Scale bar 20μm.

**Figure 4-Supplement 1.**
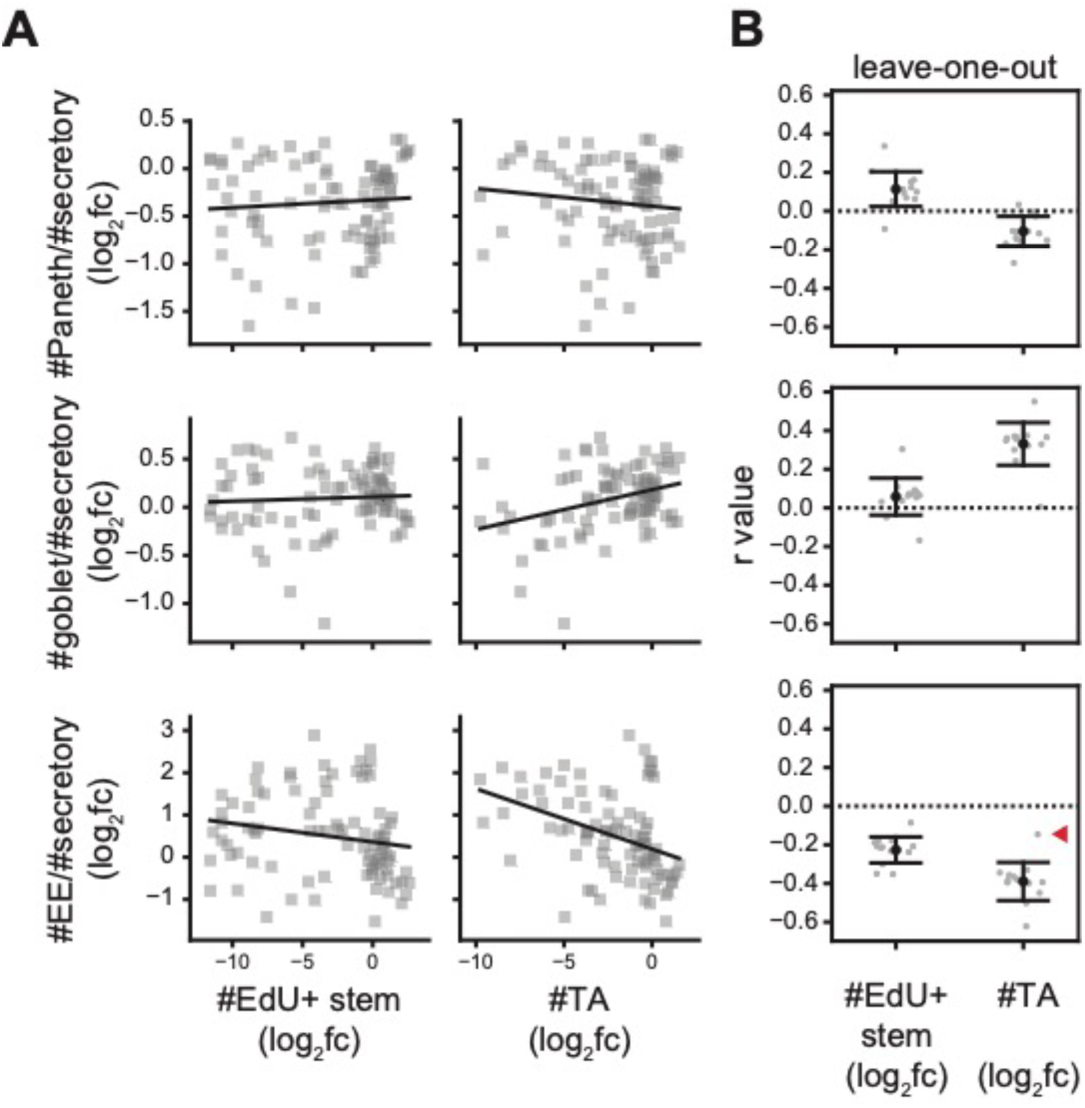
The correlation between TA cell numbers and EE cell prevalence is driven by EGFR-i. (**A**) Numbers of transit-amplifying (TA) cells correlate with EE cell fractions. Perturbation effects (log_2_fc) are plotted pairwise for each feature. (**B**) Each of 13 perturbations was sequentially dropped from the dataset and correlation coefficient (r value) calculated. Red arrowhead indicates loss of correlation between #TA cells and #EE/#secretory after dropping EGFR-i.

**Figure 4-Supplement 2.**
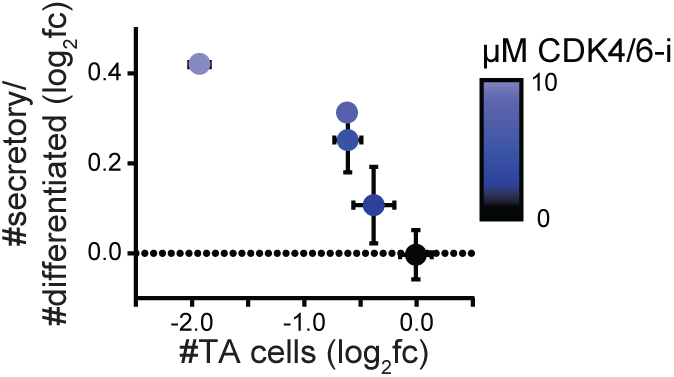
Dose-dependent increase in secretory prevalence upon CDK4/6 inhibition. Increasing amounts of CDK4/6 inhibitor were applied (concentrations indicated by color bar) to enteroid monolayers for 48 hours. Changes in the TA cells (#TA cells log_2_fc) and secretory cell prevalence amongst differentiated cells (#secretory/#differentiated log_2_fc) were quantified. The color of each point indicates the concentration of CDK4/6 inhibitor applied. n=3 wells.

**Figure 4-Supplement 3.**
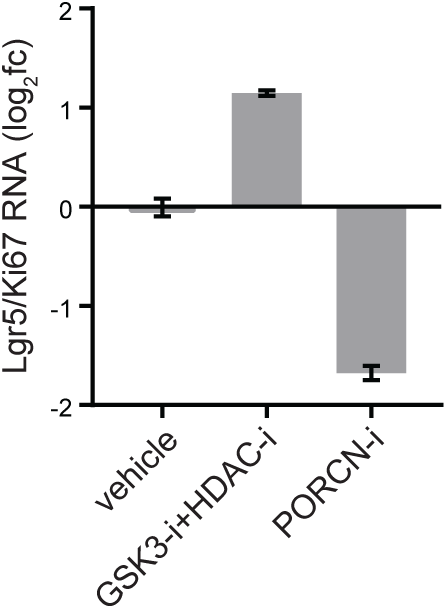
Confirming enrichment of stem versus proliferating cells in 3D organoids treated with GSK3-i+HDAC-i or PORCN-i. 3D organoids were treated with GSK3-i+HDAC-i for 2 days or PORCN-i for 1 day and relative abundance of stem (Lgr5) and proliferating (Ki67) cell RNA was measured by qRT-PCR. GSK3-i+HDAC-i increases stemness relative to proliferation and PORCN-i decreases stemness relative to proliferation. n=3 wells.

**Figure 4-Supplement 4.**
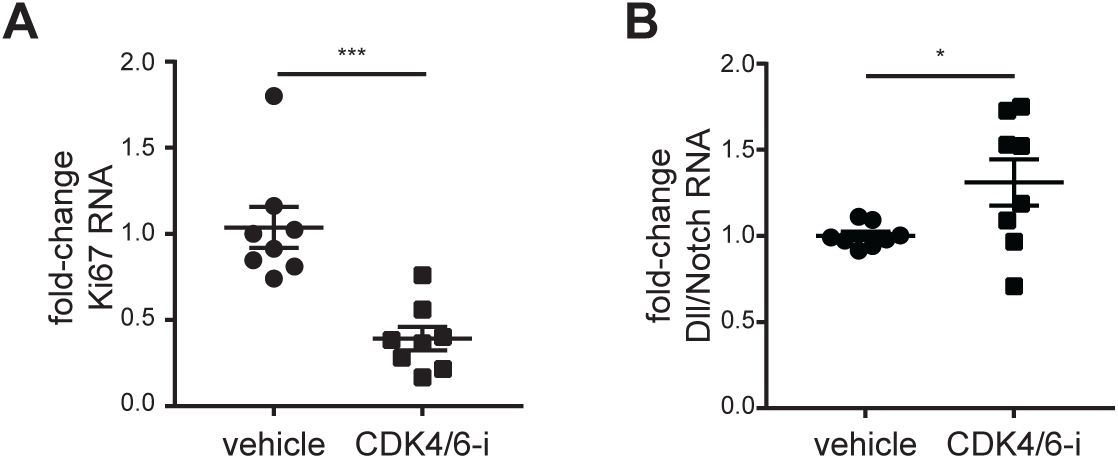
Quantifying changes in proliferation and secretory prevalence in CDK4/6-i-treated mice. Small intestinal epithelial crypts were harvested from mice treated with vehicle or the CDK4/6-i palbociclib. (**A**) To quantify changes in proliferation, Ki67 RNA was measured by qRT-PCR. CDK4/6-i decreases Ki67 expression (p=0.0003). (**B**) To quantify changes in the ratio of secretory:absorptive cell-types, Dll (Dll1+Dll4) RNA and Notch (Notch1+Notch2) RNA was measured by qRT-PCR. CDK4/6-i increases the Dll/Notch ratio, indicating an increase in secretory cell-types (p=0.039). n=8 mice/group.

**Figure 5-Supplement 1.**
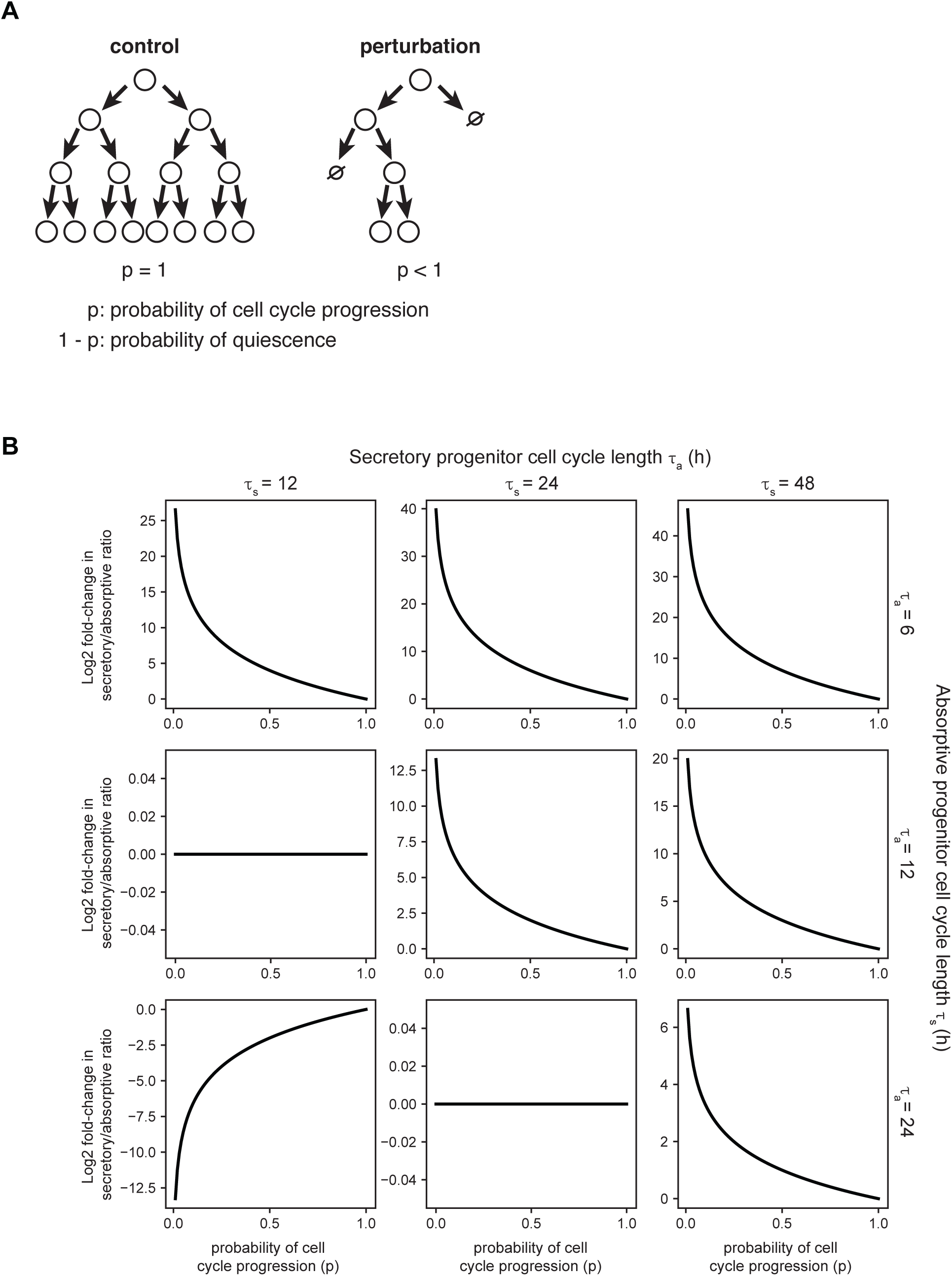
Cell cycle modulators affect the secretory:absorptive ratio due to differential amplification of secretory and absorptive progenitors. (**A**) Diagram of the effect of altering p, a parameter setting the probability of progenitor cell cycle progression. (**B**) Exponential growth model supports the hypothesis that secretory progenitors divide fewer times than absorptive progenitors. The parameters τ_a_ and τ_s_ describe how often secretory and absorptive progenitors divide. Only when secretory progenitors have longer cell cycle lengths (τ_s_>τ_a_) and thus undergo fewer divisions than absorptive progenitors during the 48-hour simulation period does decreasing the probability of cell cycle progression increase the secretory:absorptive ratio, and vice versa.

